# A practical guide to methods controlling false discoveries in computational biology

**DOI:** 10.1101/458786

**Authors:** Keegan Korthauer, Patrick K Kimes, Claire Duvallet, Alejandro Reyes, Ayshwarya Subramanian, Mingxiang Teng, Chinmay Shukla, Eric J Alm, Stephanie C Hicks

## Abstract

**Background:** In high-throughput studies, hundreds to millions of hypotheses are typically tested. Statistical methods that control the false discovery rate (FDR) have emerged as popular and powerful tools for error rate control. While classic FDR methods use only *p*-values as input, more modern FDR methods have been shown to increase power by incorporating complementary information as “informative covariates” to prioritize, weight, and group hypotheses. However, there is currently no consensus on how the modern methods compare to one another. We investigated the accuracy, applicability, and ease of use of two classic and six modern FDR-controlling methods by performing a systematic benchmark comparison using simulation studies as well as six case studies in computational biology

**Results:** Methods that incorporate informative covariates were modestly more powerful than classic approaches, and did not underperform classic approaches, even when the covariate was completely uninformative. The majority of methods were successful at controlling the FDR, with the exception of two modern methods under certain settings. Furthermore, we found the improvement of the modern FDR methods over the classic methods increased with the informativeness of the covariate, total number of hypothesis tests, and proportion of truly non-null hypotheses.

**Conclusions:** Modern FDR methods that use an informative covariate provide advantages over classic FDR-controlling procedures, with the relative gain dependent on the application and informativeness of available covariates. We present our findings as a practical guide and provide recommendations to aid researchers in their choice of methods to correct for false discoveries.

## Background

When multiple hypotheses are simultaneously tested, an adjustment for the multiplicity of tests is often necessary to restrict the total number of false discoveries. The use of such adjustments for multiple testing have become standard in areas such as genomics [1, 2], neuroimaging [3], proteomics [4], psychology [5, 6], and economics [7]. Most classically, methods which control the family-wise error rate (FWER), or probability of at least one false discovery, have been developed and used to correct for multiple testing. These include the Bonferroni correction [8, 9] and other approaches [10–12]. Despite their popularity, FWER-controlling methods are often highly conservative, controlling the probability of any false positives (Type I errors) at the cost of greatly reduced power to detect true positives. The trade-off of Type I errors and power has become exacerbated in the analysis of data from high-throughput experiments, where the number of tests being considered can range from several thousand to several million.

The false discovery rate (FDR), or expected proportion of discoveries which are falsely rejected [13], was more recently proposed as an alternative metric to the FWER in multiple testing control. This metric has been shown to have greater power to detect true positives, while still controlling the proportion of Type I errors at a specified level [13, 21]. In high-throughput biological experiments where investigators are willing to accept a small fraction of false positives to substantially increase the total number of discoveries, the FDR is often more appropriate and useful [22]. The Benjamini and Hochberg step-up procedure (BH) [13, 23] was the first method proposed to control the FDR. Soon afterwards, the *q*-value was introduced as a more powerful approach to controlling the FDR (Storey’s *q*-value) [14]. We refer to the BH procedure and Storey’s *q*-value as “classic” FDR-controlling methods (Figure 1), because they can be easily computed with just a list of *p*-values using robust software [24, 25], and are arguably still the most widely used and cited methods for controlling the FDR in practice.

**Figure 1.**
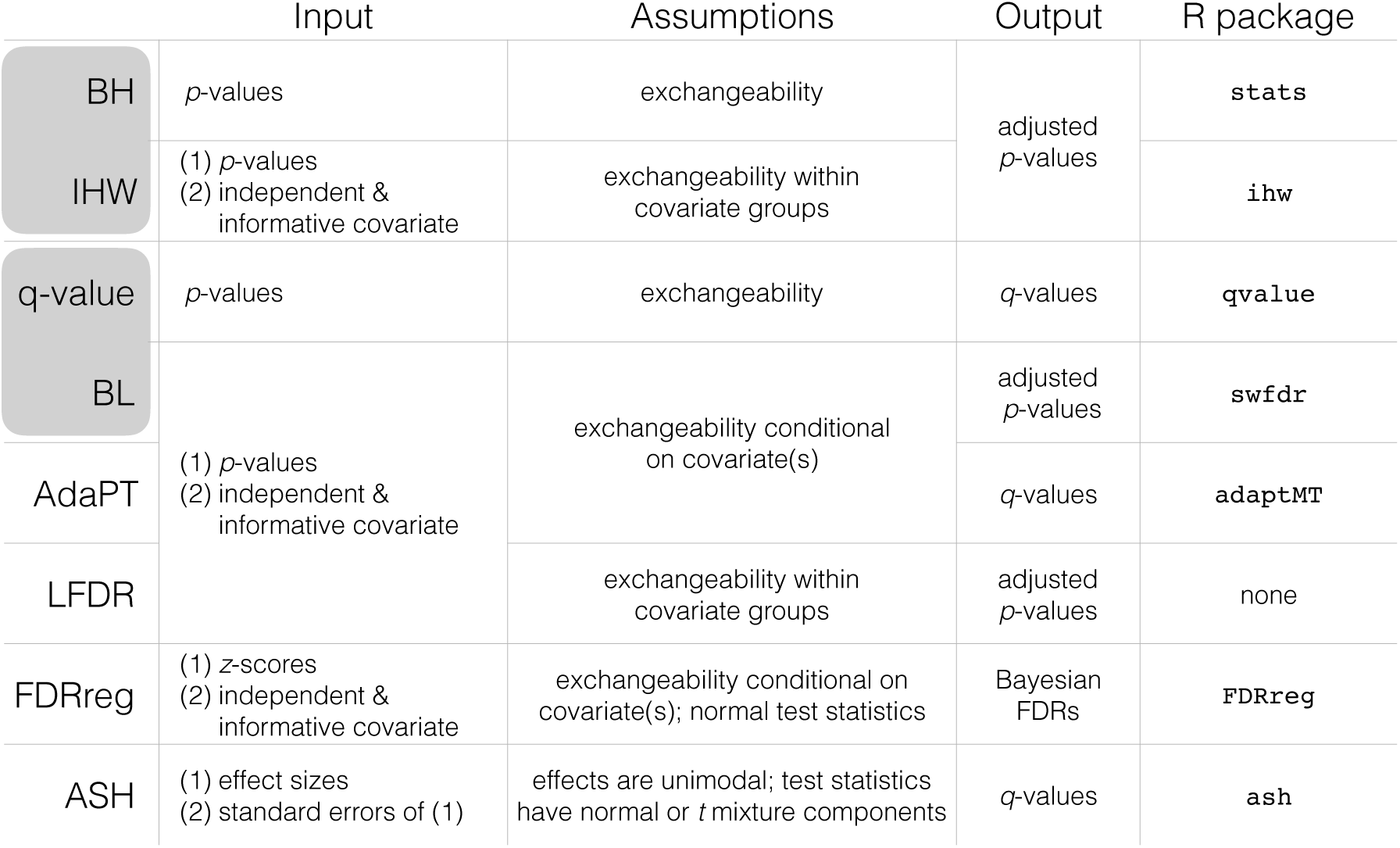
FDR-controlling methods included in the comparison. Inputs, assumptions, output, and availability (R package) of two classic [13, 14] and six modern [15–20] FDR-controlling methods. The outputs of the FDR-controlling methods vary, but they all can be used for the purpose of controlling the FDR. Pairs of classic and modern methods are highlighted in grey if the modern method is an extension of the classic method.

While the BH procedure and Storey’s *q*-value often provide a substantial increase in discoveries over methods that control the FWER, they were developed under the assumption that all tests are exchangeable, and therefore, that the power to detect discoveries is equally likely among all tests. However, individual tests or groups of tests often differ in statistical properties, such as their level of precision or underlying biology, which can lead to certain tests having greater power than others [15, 18]. For example, in a genome-wide association study (GWAS) meta analysis where samples are pooled across studies, the loci-specific sample sizes can be informative of the differing signal-to-noise ratio across loci [16]. Additionally, in an expression quantitative trait loci (eQTL) study, tests between polymorphisms and genes in *cis* are known a *priori* to be more likely to be significant than those in *trans* [15].

Recently, a new class of methods that control the FDR (Figure 1, Additional file 1: Table S1) have been proposed to exploit this variability across tests by combining the standard input (*p*-values or test statistics) [13, 14, 26] with a second piece information, referred to as an “informative covariate” [15–19, 27]. Intuitively, if a covariate is informative of each test’s power or prior probability of being non-null, it can be used to prioritize individual or groups of tests to increase the overall power across the entire experiment [15]. To guarantee FDR control, the covariate must also be independent of the *p*-values under the null hypothesis. In a similar vein, other approaches have been proposed using two alternative pieces of information, namely effect sizes and their standard errors [20], to adaptively control the FDR. These modern FDR-controlling methods allow researchers to leverage additional information or metadata and are particularly well suited for biological studies.

However, due to their recent and concurrent development, comparisons between these modern FDR methods have been limited, and the demonstration of each method’s applicability and utility on real biological problems is highly variable. Furthermore, each method requires varying sets of input data and relies on differing sets of methodological assumptions. As a result, the answer to the simple question of which methods *can,* let alone *should,* be used for a particular analysis is often unclear.

To bridge the gap between methods and application, we performed a systematic benchmark comparison of two classic and six modern FDR-controlling methods. Specifically, we compared the classic BH approach [13] and Storey’s *q*-value [14] with several modern FDR-controlling methods, including the conditional local FDR (LFDR) [17], FDR regression (FDRreg) [19], Independent Hypothesis Weighting (IHW) [15], Adaptive Shrinkage (ASH) [20], Boca and Leek’s FDR regression (BL) [16], and Adaptive p-value Thresholding (AdaPT) [18] (Figure 1). Throughout, we use lowercase when referring to the specific, typically default, implementation of each method detailed in the “Methods” section. Both the theoretical and empirical null Empirical Bayes implementations of FDRreg were compared, referred to as “fdrreg-t” and “fdrreg-e”, respectively. AdaPT was compared using the default logistic-Gamma generalized linear model option and is referenced as “adapt-glm”. The *q*-values returned by ASH were used for comparison, and are referred to as “ashq”.

Of the modern FDR-controlling methods included in our comparison, IHW, BL, AdaPT, and LFDR can be applied generally to any multiple testing problem with *p*-values and an informative covariate satisfying a minimal set of assumptions (Figure 1, Additional file 1: Table S1). In contrast, FDRreg is restricted to multiple testing problems where normal test statistics, expressed as *z*-scores, are available. Most unlike the other modern methods, ASH requires specifying effect sizes and standard errors separately for normal or *t*-distributed test statistics, and cannot be used with more general informative covariates. Furthermore, ASH requires that the true (unobserved) effect sizes across all tests are unimodal, i.e. that most non-null effect sizes are small and near zero. While this may be a reasonable assumption in settings where most non-null effects are believed to be small and larger effects are rare, it might not necessarily be true for all datasets and applications. While it is not possible to confirm whether the assumption is true, it is simple to check whether the assumption is blatantly violated, i.e. if the distribution of all observed effect sizes shows clear multimodality.

While both the BH procedure and Storey’s *q*-value serve as reference points for evaluating the modern FDR-controlling methods, in Figure 1 (and in Additional file 1: Table S1) we highlight two pairs of modern and classic methods with a special relationship: IHW with the BH procedure and BL with Storey’s *q*-value. In the case that a completely uninformative covariate is used, these modern methods have the attractive property of reducing to their classic counterparts, subject to some estimation error. Therefore, when instructive, direct comparisons are also made between IHW and the BH procedure, and similarly between BL and Storey’s *q*-value.

In this paper, we first evaluate the performance and validity of these methods using simulated data and *in silico* RNA-seq spike-in datasets. Then, we investigate the applicability of these methods to multiple testing problems in computational biology through a series of six case studies, including: differential expression testing in bulk RNA-seq, differential expression testing in single-cell RNA-seq, differential abundance testing and correlation analysis in 16S microbiome data, differential binding testing in ChIP-seq, genome-wide association testing, and gene set analysis. Combining these results with insights from our simulation studies and *in silico* experiments, we provide a key set of recommendations to aid investigators looking to take advantage of advances in multiple-testing correction in future studies.

## Results

Although fdrreg-e was included in the benchmarking study, we exclude it from the main results presentation due to its unstable and inferior performance to its counterpart fdrreg-t. For detailed results including fdrreg-e, refer to Additional file 1.

### False Discovery Rate control

The specificity of the FDR-controlling methods was evaluated using three approaches. First, a series of RNA-seq differential expression studies were performed on yeast *in silico* spike-in datasets generated by randomly selecting two sets of five and ten samples each from a dataset of 48 biological replicates in a single condition [29] and adding differential signal to a subset of genes to define “true positives”. This was carried out for a variety of settings of non-null effect size distributions, proportions of null hypotheses, and informativeness of covariates (Additional file 1: Table S2). Second, a similar differential expression study was performed using RNA-seq data simulated with the `polyester` R/Bioconductor package [28]. Finally, an extensive simulation study was carried out across a range of test statistic distributions, non-null effect size distributions, proportions of null hypotheses, informative covariates, and numbers of tests to explore a wider range of multiple testing scenarios (Additional file 1: Table S3).

All experiments and simulations were replicated 100 times. Performance metrics are reported as the mean and standard error across replications. In all analyses, covariate-aware modern FDR-controlling methods, including adapt-glm, bl, fdrreg-t, ihw, and lfdr, were run twice, once with an informative covariate and again with an uninformative random covariate.

While the notion of an informative covariate was loosely introduced above, for our *in silico* experiments and simulations, we concretely define “informative covariates” by introducing a dependence between the proportion of hypotheses that are null and the value of the covariate. A *strongly informative covariate* in our simulations is one where certain values of the covariate are highly enriched for truly non-null tests, and a *weakly informative covariate* is one where certain values are only moderately enriched for non-null tests. In contrast, an *uninformative covariate* is not enriched for null or non-null hypotheses for any values. We restrict the concepts of weakly and strongly informative covariates in our analysis to the dependence between the covariate and the null proportion described above. No other dependence is introduced between the covariate and the test statistics in our simulations and *in silico* experiments.

#### Modern methods do not always control the FDR

Across *in silico* experiments and simulation settings, we found that most methods adequately controlled the FDR in many situations. FDR control for a single setting of the yeast RNA-seq *in silico* experiments and Polyester count simulations is shown in Figure 2A. In these experiments, 30% of genes were differentially expressed (DE) between two groups of five samples each, with effect sizes sampled from a unimodal distribution and a strongly informative covariate. Here, all methods controlled the FDR at the target *α*-level, with the exception of ashq and lfdr, which exhibited slightly inflated FDR in the `polyester` simulations. Some methods, most noticeably ihw, achieved lower FDR than others. The tradeoff between FDR, power, and classification accuracy in the *in silico* experiments is summarized in Additional file 1: Figure S1.

**Figure 2.**
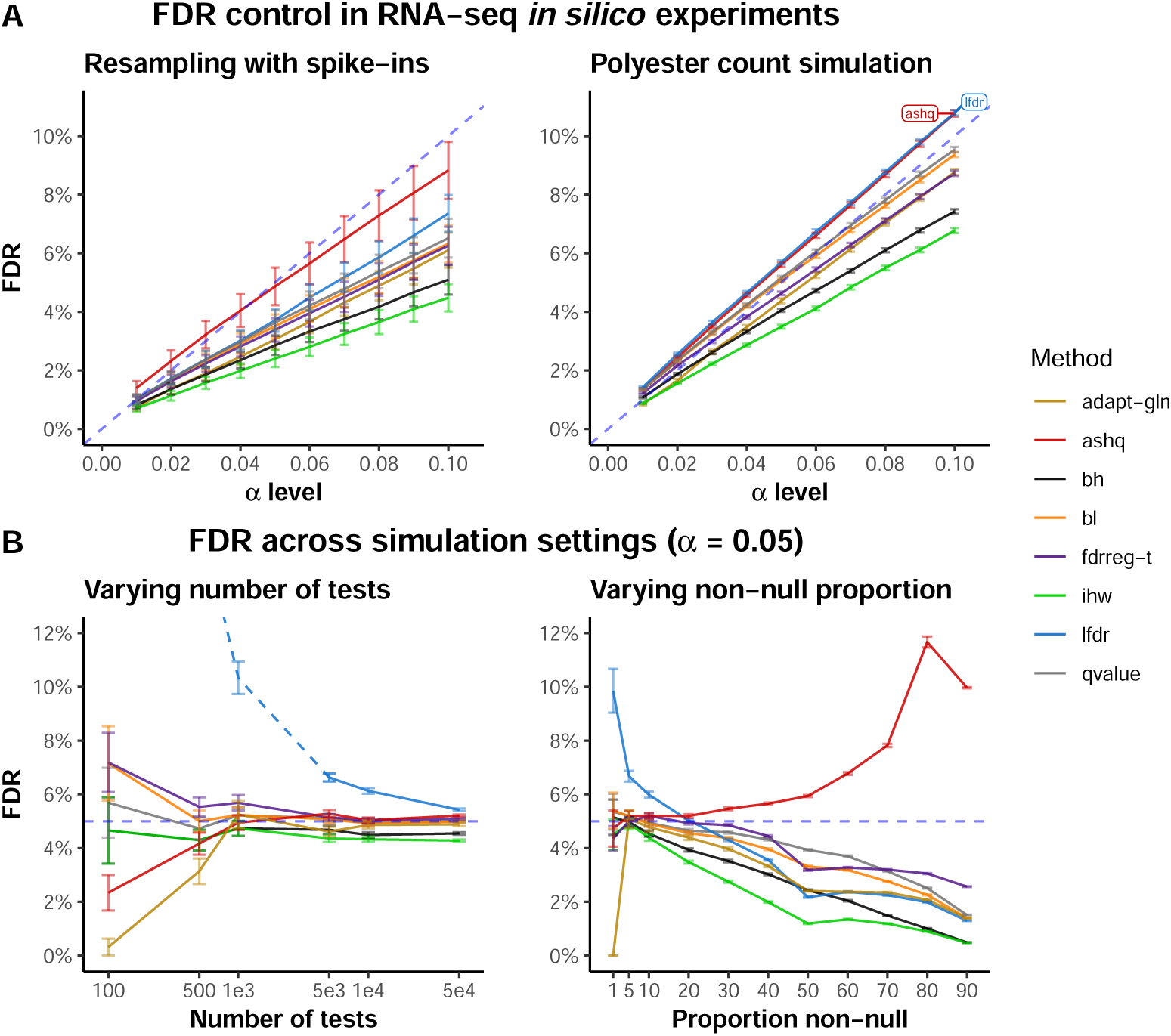
FDR control in *in silico* experiments and simulations. (A) Observed FDR (y-axis) for various *α*-level cutoffs (x-axis) in the yeast RNA-seq *in silico* resampling experiment with spiked-in differentially expressed genes (left panel) and the simulation of yeast RNA-seq counts using the `polyester` R/Bioconductor package [28]. (B) Observed FDR (y-axis) across simulation settings at *α*-level of 0.05. The left panel displays FDR for increasing numbers of hypothesis tests and the right panel displays FDR for increasing proportions of non-null hypotheses. Note that the lfdr method is displayed as a dotted line when the number of tests per bin falls below 200 (where the number of bins is fixed at 20), as `fdrtools` generates a warning in this case that the estimation may be unreliable.

The settings of the *in silico* experiments were varied to also consider a lower proportion of DE genes (7.5%), bimodal effect size distribution, and a weakly informative covariate in addition to the uninformative random covariate run with all covariate-aware methods (Additional file 1: Table S2). The FDR for covariate-aware methods was not sensitive to covariate informativeness, with nearly identical proportions of false discoveries using weakly and strongly informative covariates. However, we found that with the bimodal effect size distribution and smaller proportion of non-null hypotheses, a subset of methods including ashq and lfdr failed to control the FDR at the nominal FDR cutoff (*α*), leading to an inflated rate of false discoveries (Additional file 1: Figure S2).

Similar trends were observed across simulation studies, where conditions were varied analogous to the yeast experiments to consider a wider range of scenarios. While most methods were consistently conservative or achieved an accurate target FDR, some methods clearly failed to control the FDR under certain settings.

#### lfdr and fdrreg-t do not control FDR with few tests

Since modern FDR-controlling methods must estimate the covariate dependence from the set of hypotheses, the effectiveness of these methods can depend on having a sufficiently large number of tests. We performed a series of simulations to assess the sensitivity of the covariate-aware methods to the total number of hypotheses. We observed that lfdr exhibited substantially inflated FDR when applied to 10,000 or fewer tests (Figure 2B, left panel). This result could be due to our implementation of LFDR, which groups hypotheses into 20 groups regardless of the total number of tests, and suggests that the performance of lfdr improves when the numbers of tests per bin increases. We also observed that fdrreg-t showed slightly inflated FDR with 1,000 or fewer tests.

#### lfdr and ashq do not control FDR for extreme proportions of non-null tests

The proportion of non-null tests is typically unknown, but can vary dramatically between data sets. While most simulation settings were performed with 10% non-null tests, to cover a range of scenarios a series of settings covering non-null proportions between 0% and 95% were also considered. The yeast *in silico* experiments included settings of 0%, 7.5% and 30% non-null tests.

Most methods demonstrated the same general trend in simulation, where the FDR of most methods was controlled at the target *α*-level, and decreased as the proportion of non-null hypotheses increased (Figure 2B, right panel). However, we also found that the ability of some methods to control FDR was sensitive to the proportion of tests that were non-null. Specifically, lfdr exhibited inflated FDR when the proportion of non-null tests was low (less than 20%). Likewise, ashq exhibited inflated FDR when the proportion of non-null tests was high (greater than 20%).

Similarly, ash and lfdr failed to control FDR in the *in silico* yeast experiments when the proportion of non-nulls was 7.5% compared to 30% (Additional file 1: Figure S2). We also note that for a sample size of 5 per group, several methods exhibited inflated FDR in the extreme setting when the non-null proportion of hypotheses was 0%, where FDR reduces to FWER. However, although the proportion of replications with at least one false positive was greater than the target, the average proportion of tests rejected was very small (Additional file 2). Since the *in silico* experiments were generated by splitting biological replicates into two groups, it is possible that unmeasured biological differences exist between them.

### Power

In addition to FDR, we also evaluated sensitivity of the FDR-controlling methods using the same *in silico* experiment and simulation framework described above.

#### Modern methods are modestly more powerful

We found that in general modern FDR methods led to a modestly higher true positive rate (TPR), or power, in the yeast *in silico* RNA-seq experiments and `polyester` simulations (Figure 3A). This was also true when using a weakly informative rather than a strongly informative covariate (Additional file 1: Figure S2B). Much of the gain with modern methods, most apparent with lfdr and ashq, was found in genes with small to moderate effect sizes (Additional file 1: Figure S1D). While the majority of discoveries were common among all or most methods, there were several smaller sets of rejections that were unique to a small number of methods (Additional file 1: Figure S1E).

**Figure 3.**
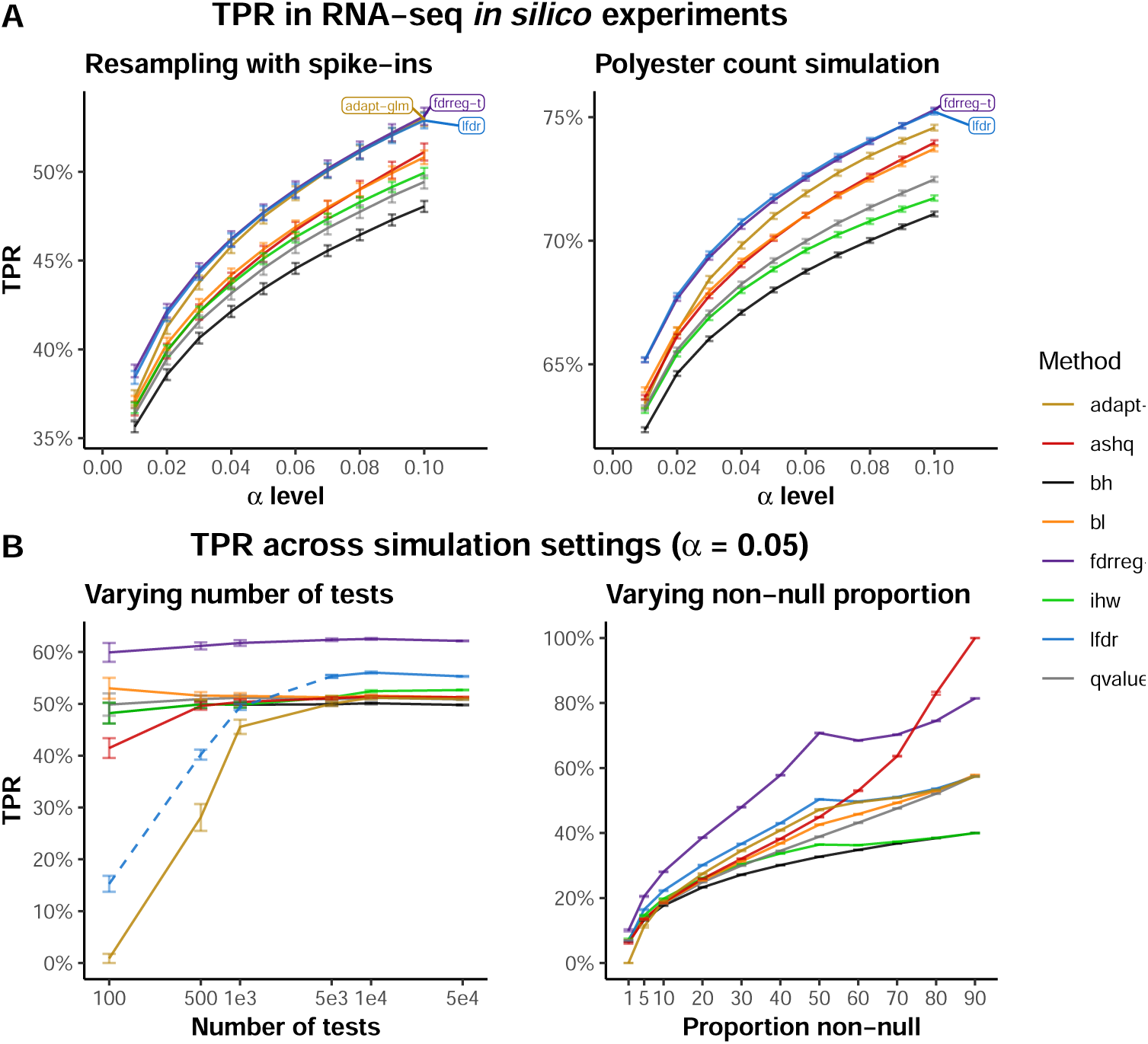
Power in *in silico* experiments and simulations. (A) True Positive Rate (y-axis) for increasing *α*-level cutoffs (x-axis) in the yeast RNA-seq *in silico* resampling experiment with spiked-in differentially expressed genes (left panel) and the simulation of yeast RNA-seq counts using the `polyester` R/Bioconductor package [28]. (B) True Positive Rate (y-axis) across simulation settings at *α*-level of 0.05. The left panel displays increasing numbers of hypothesis tests and the right panel displays increasing proportions of non-null hypotheses. Note that the lfdr method is displayed as a dotted line when the number of tests per bin falls below 200 (where the number of bins is fixed at 20), as `fdrtools` generates a warning in this case that the estimation may be unreliable.

Again, higher power was similarly observed for most modern FDR methods over classic methods across simulation settings (Figure 3B, Additional file 1: Figures S3, S4, S5, S6, S7). The increase in TPR was generally modest for all methods, as in the yeast experiments, with the exception of fdrreg-t which showed substantial improvement in TPR over modern methods in several simulation settings (Additional file 1: Figure S3B, D, and F).

#### Power of modern methods is sensitive to covariate informativeness

Comparing across yeast experiments using weakly and strongly informative covariates, we found that the TPR was higher for strongly informative covariates compared to weakly informative covariates (Additional file 1: Figure S2). To further quantify the impact of covariate informativeness, a series of simulations was performed using covariates of varying informativeness. A rough scale of 0-100 was used to describe the informativeness of the covariate, with larger values of informativeness corresponding to greater power of the covariate to distinguish null and non-null tests (Additional file 1: Figure S8). Echoing the results of the yeast experiments, the gain in TPR of covariate-aware methods over other methods also increased with informativeness in the simulation studies (Additional file 1: Figure S3B). This gain tended to be larger for some methods (fdrreg-t, lfdr, and adapt-glm) than for others (ihw and bl).

Additionally, simulations were performed across four different dependencies between the covariate and the null proportion. The covariates were named *step*, *cosine*, *sine*, and *cubic* for the shape of their dependence. We found that in general, the above results were relatively robust to the functional form of the dependence (Additional file 1: Figure S4A-C). However, the gain in TPR varied across informative covariates, with the smallest gains observed in the *step* covariate setting across all covariate-aware methods, likely attributable to the lower informativeness of the covariate relative to the other settings. The gain in TPR also varied more for some methods than others. In the *cosine* covariate, where the dependence between the covariate and null proportion was strongly non-monotone, bl showed no gain over the classic *q*-value. As bl attempts to model the covariate-null proportion dependence using logistic regression, a monotone function, the method was unable to capture the true relationship. A small, but noticeable increase in TPR was observed in the remaining settings, where the covariate dependence was monotone. In contrast, covariate-aware methods with more flexible modeling approaches either based on binning (ihw, lfdr) or spline expansion (fdrreg-t), were generally more consistent across covariates.

#### Including an uninformative covariate is not harmful

A reasonable concern, closely related to weakly and strongly informative covariates, is whether an uninformative covariate could mislead methods such as ihw, bl, fdrreg, lfdr, or adapt-glm. Across settings of the yeast *in silico* experiments and simulations, we observed that with the use of a completely uninformative covariate, modern FDR methods generally had lower power (and higher FDR) than with an informative covariate (Additional file 1: Figures S4D-E, S5D-E, S6D-E, S9, and S10B). However, while modern FDR methods were modestly more powerful than classic approaches when using an informative covariate, they did not underperform classic approaches with a completely uninformative covariate (Additional file 1: Figure S3A-B).

A notable exception was adapt-glm, which suffered from lower power with the inclusion of a weakly informative covariate than with the uninformative covariate, likely due to overfitting (Additional file 1: Figures S3B and S4E). In estimating the dependence between the covariate and null proportion, adapt-glm includes a step of model selection. Based on feedback from the method authors, we considered modifying the default adaplt-glm parameters by including a null dependence as one of the model choices, allowing the method to ignore the dependence when it cannot be properly estimated. When applied to the weakly informative *step* covariate setting, this resulted in improved performance with the method no longer suffering from lower power with the inclusion of the weakly informative covariate (Additional file 1: Figure S11). However, since this procedure was not used in [18] and is not currently mentioned in the software documentation, we have excluded it from our primary analyses. The authors responded positively to the recommendation of documenting this procedure in future releases of the package.

#### lfdr and adapt-glm are sensitive to the number of tests

We found that the power of some methods was more sensitive to the number of hypothesis tests in the simulation studies than others. Specifically, lfdr and adapt-glm performed poorly in terms of TPR when there were fewer than 1000 tests (Figure 3B, left panel). The lfdr result may again be due to our implementation, as described above. We also note that the improvement in TPR of ihw over bh was not apparent unless there were at least several thousand tests (Additional file 1: Figure S3D).

### Applicability

To investigate the applicability of modern methods to a variety of analyses and datasets, we used a combination of simulation settings and empirical case studies. Specifically, we evaluated performance under several different test statistic and effect size distributions in simulation. We considered normal, *t* with both 5 and 11 degrees of freedom, and *χ*^2^ test statistics with 4 degrees of freedom as in [16]. Additionally, we considered several different effect size distributions, ranging from unimodal to bimodal.

We also investigated the application of these methods to a series of six case studies in computational biology, including: differential expression testing in bulk RNA-seq, differential expression testing in single-cell RNA-seq, differential abundance testing and correlation analysis in 16S microbiome data, differential binding testing in ChIP-seq, genome-wide association testing, and gene set analysis. These results, along with a practical discussion of the selection and assessment of informative covariates, are included in the following sections.

#### ashq and fdrreg-t are sensitive to the sampling distribution of the test statistic

Many of the modern FDR-controlling methods make assumptions regarding a valid distribution of *p*-values. However, some methods also make assumptions about the distribution of the test statistic or effect size. Specifically, FDRreg and ASH both assume that test statistics are normally distributed [19, 20]. However, ASH is also described as being applicable to *t*-distributed statistics, although, currently only based on a rough approximation [20]. The option to specify the degrees of freedom for *t*-distributed statistics based on this approximation was used for the ashq implementation in the *t*-distributed simulations. The sensitivity of these methods along with the others to changes in the underlying distributions of the test statistics was investigated through simulations across four distributions: normal, *t* with 11 and 5 degrees of freedom, and *χ*^2^ with 4 degrees of freedom. These simulation results are shown in Figure 4A and Additional file 1: Figure S5. Since the assumptions for both FDRreg and ASH are strongly violated with *χ*^2^ test statistics, these methods were not applied in this setting.

**Figure 4.**
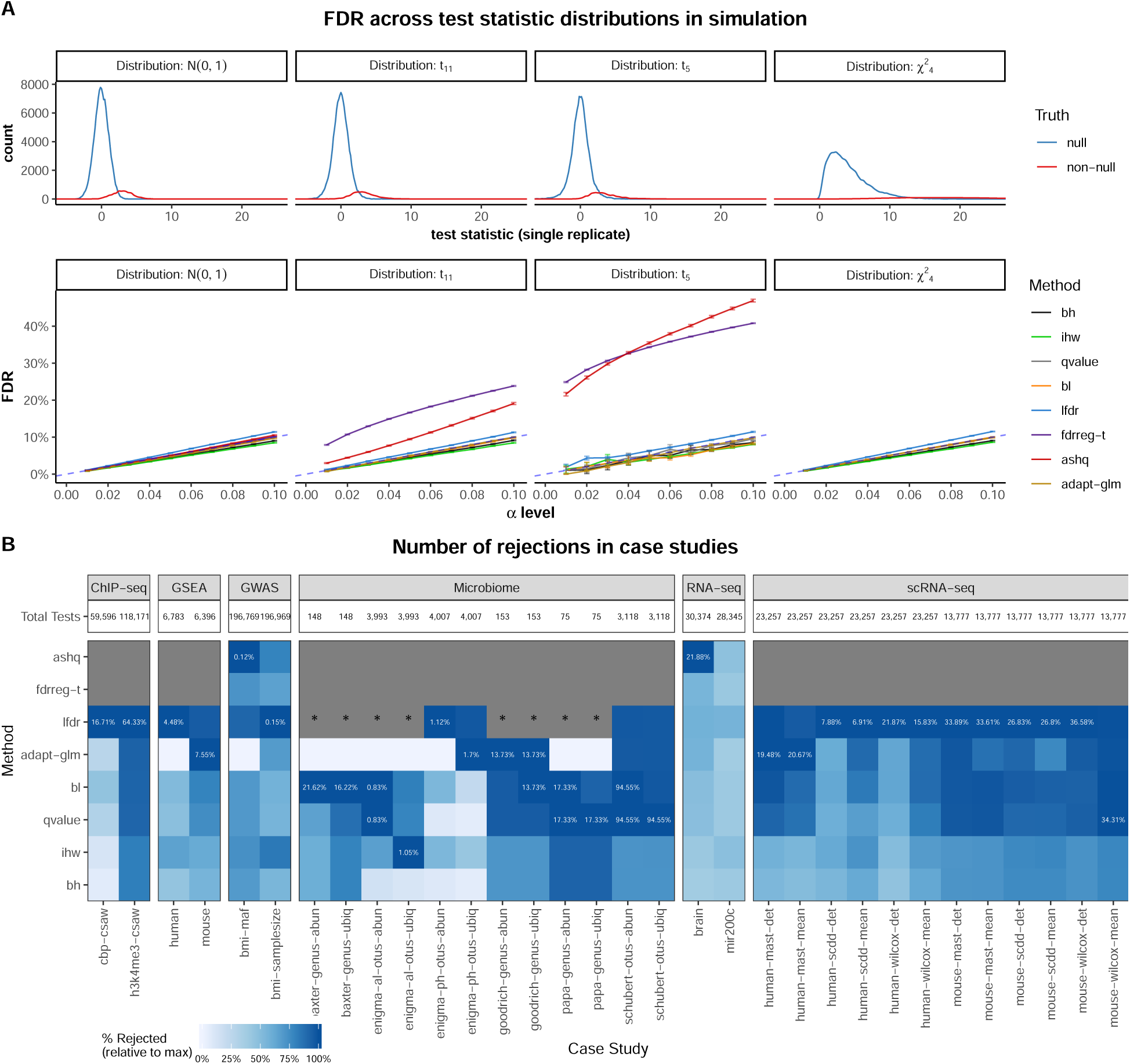
Applicability of benchmarked methods to various test statistics and case study datasets. (A) The top panel displays various null and non-null test statistic distributions used in simulations, with the corresponding observed FDR shown in the bottom panel. Note that although fdrreg-t requires normally distributed test statistics, it is included in the evaluation of *t*_11_ and *t*_5_ statistics to illustrate the effect of a heavy tailed distribution. In addition, neither ashq nor fdrreg-t are evaluated on *χ*_2_ statistics, as they violate assumptions of the method. (B) Proportion of maximum rejections (color) for each dataset and informative covariate (column, grouped by case study) and FDR correction method (row). In each column, the maximum proportion of rejections out of the total possible number of comparisons is displayed. Where methods ashq and fdrreg-t could not be applied to a case study to due violation of assumptions, the cell is colored in grey. Where the method lfdr was not applied due to practical limitations on the number of tests, the cell is colored grey and marked with “*”. The informative covariate used in each case study is listed in Table 1.For case studies with more than one covariate, the covariate is denoted in the *x*-axis labels.

We observed that FDR control for most methods, namely those which take *p*-values as input rather than *z*-scores or effect sizes (Figure 1), was not sensitive to the distribution of test statistics (Figure 4A and Additional file 1: Figure S5B-C). However, violation of the normal assumption of fdrreg-t led to inflated FDR when the test statistics were *t*-distributed, and as expected, the increase in FDR was greater for the heavier-tailed *t* distribution with fewer degrees of freedom (Figure 4A). Although it accommodates *t*-distributed test statistics, inflated FDR was also observed for ashq (both with and without specifying the correct degrees of freedom). In personal communications, the authors of [20] have acknowledged that the current procedure for *t*-distributed test statistics can be improved and are actively developing an adaptation of ashq for this case.

#### ashq is not sensitive to violation of the unimodal assumption

In addition to distributional assumptions on the test statistic, ASH assumes that the distribution of the true (unobserved) effect sizes is unimodal, referred to as the ‘unimodal assumption’. To investigate ASH’s sensitivity to the unimodal assumption, multiple distributions of the effect sizes were considered in both simulations and yeast *in silico* experiments. Both effect size distributions following the unimodal assumption of ASH and those with most non-null effects away from zero (Additional file 1: Figure S5A), were considered. While most simulations included the latter, simulations were also performed with a set of unimodal effect size distributions described in [20] (Additional file 1: Figure S6 and S7). In the yeast *in silico* experiments, two conditions were investigated - a unimodal and a bimodal case.

We also observed that even when the unimodal assumption of ASH was violated in simulation, ashq had only a slight inflation in FDR and comparable TPR to other methods (Additional file 1: Figure S6B-C). This was also observed in the yeast *in silico* experiments (Additional file 1: Figure S2).

#### Not all methods could be applied to every case study

We discovered that some methods could not be applied to some case studies due to restrictive assumptions. For example, FDRreg could only be applied if the tests under consideration yielded approximately normally-distributed statistics. As a result, FDRreg was applied to the bulk RNA-seq and GWAS studies, but not considered in any of the other case studies since the test statistics are decidedly not normal. Likewise, ASH could only be applied if both an effect size and corresponding standard error for each test was available. As a result, ASH was excluded from case studies involving tests that only output a *p*-value or test statistic, such as permutation tests or the Wilcoxon rank-sum test (Figure 4B). Further, the lfdr method was not applied to three microbiome datasets where there were fewer than 4,000 total tests (200 tests per bin).

In the *in silico* experiments and simulations described above where the underlying properties of the data are known, it is easy to verify whether the assumptions of each method are satisfied. In practice, however, some assumptions are difficult or even impossible to check. For example, while it is feasible to assess the overall unimodality of the observed effect sizes for input to ashq, it is impossible to check the ‘unimodal assumption’ for the true (unobserved) effects. For this reason, it is possible that the assumptions of ashq could be violated in some of the case studies.

#### Choice of independent covariate was application-dependent

Several covariates have been suggested for *t*-tests, rank-based tests, RNA-seq DE analysis, eQTL analysis, GWAS, and quantitative proteomics [15]. In the case studies, we selected covariates based on these suggestions, as well as our own hypotheses about covariates that could potentially contain information about the power of a test, or the prior probability of a test being non-null (Table 1). We observed that the relationship between the covariates explored in the case studies and the proportion of tests rejected was highly variable (Additional file 1: Figure S12).

**Table 1.** Independent and informative covariates used in case studies.

| Case study | Covariates found to be independent and informative |
| --- | --- |
| Microbiome | **Ubiquity**: the proportion of samples in which the feature is present. In microbiome data it is common for many features to go undetected in many samples.<br>**Mean nonzero abundance**: the average abundance of a feature among those samples in which it was detected. We note that this did not seem as informative as ubiquity in our case studies. |
| GWAS | **Minor allele frequency**: the proportion of the population which exhibits the less common allele (ranges from 0 to 0.5). Represents the rarity of a particular variant.<br>**Sample size (for meta-analyses)**: the number of samples for which the particular variant was measured. |
| Gene Set Analyses | **Gene set size**: the number of genes included in the particular set. Note that this is not independent under the null for over-representation tests, however (see Additional file 1: Supplementary Results). |
| Bulk RNA-seq | **Mean gene expression**: the average expression level (calculated from normalized read counts) for a particular gene. |
| Single-Cell RNA-seq | **Mean nonzero gene expression**: the average expression level (calculated from normalized read counts) for a particular gene, excluding zero counts.<br>**Detection rate**: the proportion of samples in which the gene is detected. In single-cell RNA-seq it is common for many genes to go undetected in many samples. |
| ChIP-seq | **Mean read depth**: the average coverage (calculated from normalized read counts) for the region<br>**Window Size**: the length of the region |

To select covariates for each case study, we visually evaluated whether each covariate was informative by examining a scatter plot of the independent covariate percentile and the *p*-value. If this contained any sort of trend such that certain values of the informative covariate were enriched for smaller *p*-values, we considered the covariate to be informative.

We also visually evaluated whether each covariate was approximately independent under the null hypothesis following the recommendations of [15]. Specifically, we examined the histogram of *p*-values stratified by small, moderate and large values of the covariate (Additional file 1: Figure S13). If the distribution of the moderate to large *p*-values appeared approximately uniform, we considered the covariate to be approximately independent of the *p*-values under the null hypothesis.

For almost all choices of the covariate, we were able to substantiate evidence for informativeness and independence. One notable exception was the set size covariate for the overrepresentation test in the gene set analysis case study. Here, we found that although the covariate appeared to be informative, it was not independent under the null hypothesis (Additional files 22-23). We observed a dependence in the global enrichment of smaller *p*-values in small gene sets. This is a direct consequence of the fact that a single DE gene represents a larger proportion of a smaller gene set than it does a larger gene set. As a result, we only show results for gene set analysis using Gene Set Enrichment Analysis (GSEA), which does not rely on selecting a subset of DE genes, but instead incorporates the rank of every gene into the evaluation of a gene set. The gene set size covariate did satisfy the independent and informative criteria for *p*-values obtained from GSEA.

### Consistency

We observed that the relative performance of modern methods differed depending on the particular scenario. To evaluate the consistency of the performance of modern methods, we summarized the variability across the different simulation studies, *in silico* experiments, and case studies.

Across all simulation studies and yeast *in silico* experiments, we quantified the overall proportion of settings of modern FDR methods achieving FDR control (Figure 5A) and the average ranking of TPR (Figure 5B). In addition, we quantified the variability across simulation settings of modern FDR methods relative to classic methods (Figure 5C-D). We also evaluated the consistency of the number of rejections in case studies both with and without informative covariates. Note that variability across case studies was not evaluated for fdrreg and ashq, as the methods were only applied to a subset of the datasets. Detailed discussion of these results is provided in the following sections.

**Figure 5.**
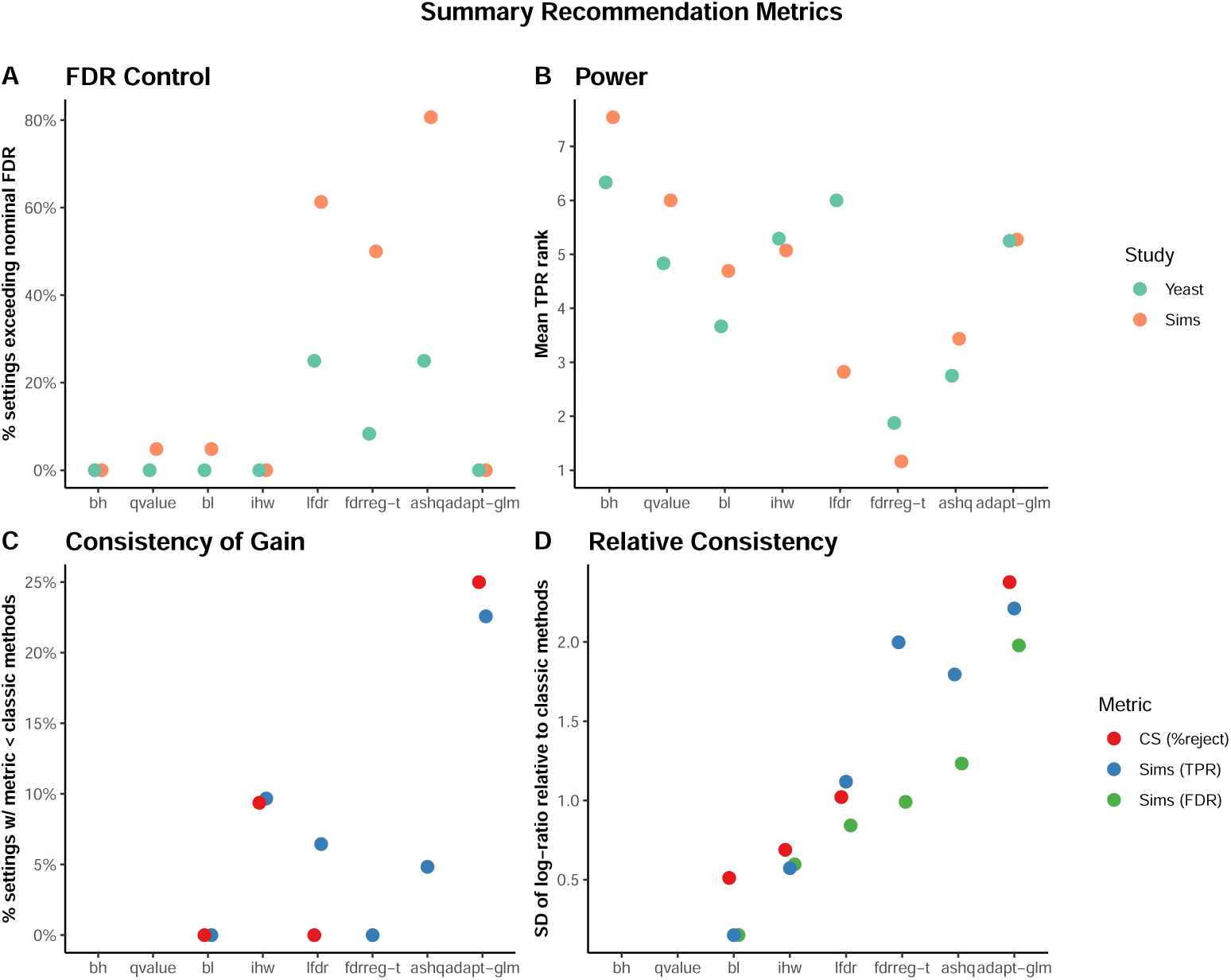
Summary metrics computed to rate methods for final recommendations. Several metrics were computed over all settings of the simulations, and yeast experiments, as well as all datasets and covariates in the case studies to evaluate the (A) FDR control, (B) power, and (C,D) consistency of the evaluated methods. In panels (A) and (B), color denotes whether the metric is computed over simulations (“Sims”) or yeast *in silico* experiments (“Yeast”). In panels (C) and (D), color denotes whether the metric is computed for TPR or FDR in the simulations (“Sims (TPR)” and “Sims (FDR)”, respectively), or for the percentage of rejections in the case studies “CS (%reject)”. In all panels, methods are shown on the *x*-axis, and methods with superior performance are those with a low value of the *y*-axis metric. Cutoffs used with the metrics shown are provided in the “Methods” section.

#### Consistency of FDR and Power

We observed that adapt-glm, ihw, and bl achieved FDR control in almost all simulation and *in silico* experiment settings (Figure 5A), and were ranked near the median of all methods in terms of TPR on average (Figure 5B). However, adapt-glm frequently resulted in lower TPR than classic methods (Figure 5C), and had the highest variability of TPR and FDR across all simulation settings (Figure 5D). Note that although ihw had lower TPR than bh and qvalue in about 10% of simulation settings (Figure 5C), this difference was usually small and the variability of ihw relative to classic methods was smaller than most other modern methods (Figure 5D).

On the other hand, fdrreg-t and ashq were consistently ranked among the top methods in terms of TPR (Figure 5B), but both failed to control FDR in more than 40% of simulation settings (Figure 5B) and exhibited higher variability of both FDR and TPR than bl and ihw (Figure 5D). lfdr showed similar performance to ashq and fdrreg-t, but was ranked more favorably in terms of TPR in simulation studies compared to *in silico* experiments (Figure 5).

#### Number of rejections highly variable in case studies

In the case studies, we found that lfdr and ashq (where applied) made the most rejections on average (Additional file 1: Figure S14), a similar trend to that observed in the yeast *in silico* simulations (Additional file 1: Figure S15). Otherwise, the relative ranking among the methods varied among datasets and covariates used in each analysis (Figure 4B).

The adapt-glm and lfdr methods had the most variable performance relative to classic methods across case studies (Figure 5D). In particular, adapt-glm rejected fewer tests than the classic methods in approximately 25% of case study datasets (Figure 5C). The performance pattern of bl was very similar to qvalue (Figure 4). Likewise, ihw exhibited similar patterns to bh. The ashq method, where applied, was usually among the methods with the most rejections, and bh consistently found among the fewest discoveries on average among all FDR-controlling methods.

#### Gain over uninformative covariates highly variable in case studies

To investigate how each method uses information from covariates and to assess performance in the case that a covariate is completely uninformative, we also included a randomly generated uninformative covariate in each case study, that was independent of the *p*-values under the null and alternative.

The average gain from using an informative covariate as compared to an uninformative covariate was usually modest, but in rare cases resulted in order of magnitude differences (Additional file 1: Figure S10A). The gain was also highly variable across case studies, covariates, and datasets. In some cases the adapt-glm and bl methods made fewer rejections using the informative covariate (Additional file 1: Figure S16).

## Discussion and Conclusions

We have presented a systematic evaluation to guide researchers in their decisions regarding methods to control for false discoveries in their own data analysis. A series of case studies and simulations were performed to investigate which methods maximize the number of discoveries while controlling FDR at the nominal *α*-level. We conclude by highlighting several key results and practical recommendations, which are summarized in Figure 6.

**Figure 6.**
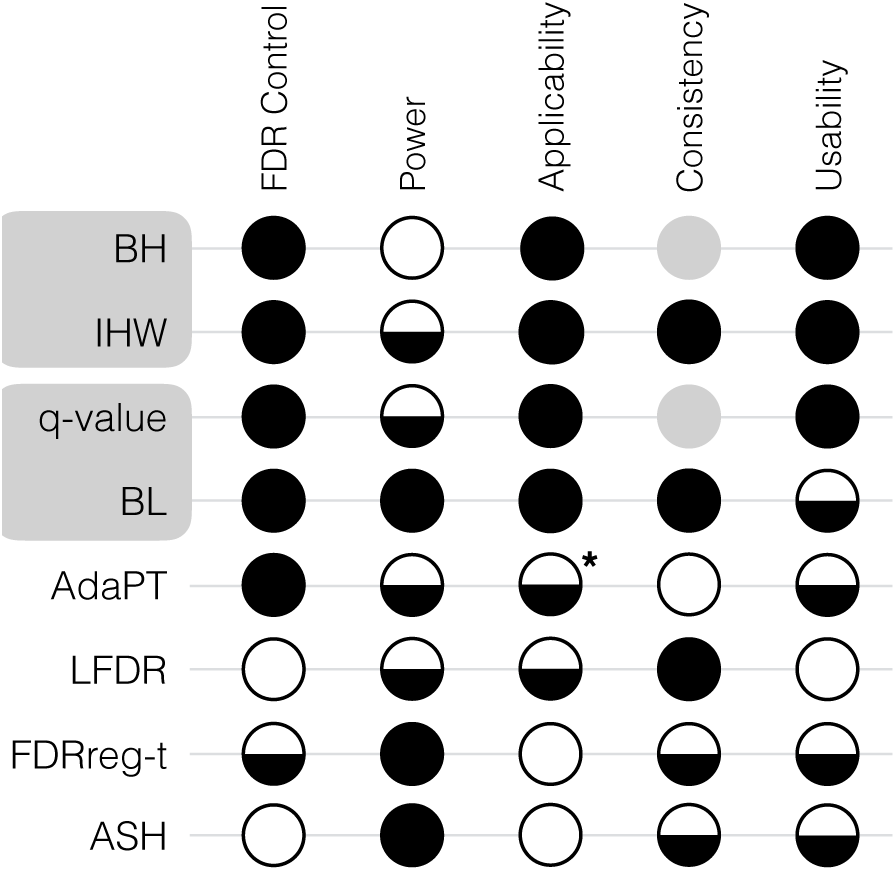
Summary of recommendations. For each method (row) and evaluation criteria (column), a filled circle denotes the method was superior, a half-filled circle denotes the method was satisfactory, and an empty circle denotes the method was unsatisfactory. Gray circles are used to denote that BH and *q*-value were not evaluated for the Consistency criteria. An asterisk is used to denote that applicability was assessed slightly differently for AdaPT. Detailed evaluation criteria are provided in the “Methods” section.

We found that modern methods for FDR control were more powerful than classic approaches, but that the gain in power was generally modest. In addition, with the exception of AdaPT, most methods that incorporate an independent covariate were not found to underperform classic approaches, even when the co-variate was completely uninformative. Because adapt-glm sometimes performed worse with the use of a co-variate, we recommend including a null model as input along with the covariate model when applying AdaPT.

Overall, we found the performance of the modern FDR methods generally improved over the classic methods as (1) the informativeness of the covariate increased, (2) the number of hypothesis tests increased, and (3) the proportion of non-null hypotheses increased. Although it is not possible to assess (1) and (3) in practice, most methods still controlled FDR when the covariate was weakly informative and the proportion of non-nulls was high.

Across our simulation and case study evaluations, we found that IHW and BL generally had the most consistent gains in TPR over classic methods, while still controlling the FDR (Figure 6). While the TPR of BL was often higher than IHW, we note that the gain in power of BL relative to IHW should be interpreted in light of any gain in power of *q*-value to BH, due to the special relationship between these pairs of methods. Specifically, IHW and BL reduce to BH and *q*-value, respectively, when the covariate is uninformative. The power of IHW was generally superior to BH when the covariate was sufficiently informative, but almost identical to BH when the covariate was not informative enough or when there were only a few thousand tests. Likewise, the power of BL was generally superior to Storey’s *q*-value when the covariate was sufficiently informative and had a monotonic relationship with the probability of a test being non-null.

We also found that although the majority of methods performed similarly in controlling the FDR, some methods were not able to control FDR at the desired level under certain settings. This occurred for empirical FDRreg when the proportion of non-nulls was near 50%, LFDR when there were fewer than 5000 tests, and ASH when the test statistic was *t*-distributed.

We have provided several useful examples of how to use an informative covariate in biological case studies. When choosing a covariate for a particular analysis, it is important to evaluate whether it is both informative and independent under the null hypothesis. In other words, while the covariate should be informative of whether a test is truly positive, if a test is truly negative, knowledge of the covariate should not alter the validity of the *p*-value or test statistic. Violation of this condition can lead to loss of Type I error control and an inflated rate of false discoveries [30]. To avoid these pitfalls, we recommend using previously proposed visual diagnostics to check both the informativeness and independence of the selected covariate [15].

We also note that although in this study we only considered a single (univariate) covariate in the simulations and case studies, some of the modern methods are able to incorporate multiple covariates. BL, AdaPT, and FDRreg can all accommodate an arbitrary set of covariates through the specification of a design matrix. In particular, AdaPT is well-suited to high-dimensional problems, as it provides an implementation that uses *L*_1_-penalized Generalized Linear Models for feature selection. Further investigation is needed in the selection of multiple covariates and the potential gain in performance over using a single co-variate.

Finally, we rank IHW, BH, and Storey’s *q*-value as superior in terms of user-friendliness and documentation, critical for lasting use and impact in the community. All methods were implemented and evaluated in R. With the exception of LFDR and the latest version of FDRreg, methods were easily accessible from packages in CRAN or Bioconductor, the primary repositories for R packages. Implementing LFDR and installing FDRreg both required additional work (see the “Methods” section). In their implementations, most methods provide direct measures, such as adjusted *p*-values or *q*-values, as outputs directly to users. In contrast, bl provides null hypothesis weights which must be manually applied to BH-adjusted p-values by the user to control FDR. In addition, bl, adapt-glm and fdrreg, all require specifying a functional relationship between the covariate and null proportion in the form of a model or formula. While this provides the user significant flexibility, it can also be unintuitive for researchers not familiar with the underlying modeling frameworks of these methods. A benchmarking experiment to determine reasonable default values for parameters to improve the user-friendliness of these methods is left as future work.

## Methods

### Assessing assumptions

ASH and FDRreg differ substantially from the remaining methods, and care should be taken to verify that the appropriate inputs are available and that the underlying assumptions are indeed valid. Based on these criteria, both methods were excluded from most case studies and several simulation settings considered in this benchmark. A more detailed discussion of these assumptions is included below. For FDRreg and adapt-glm, the informative covariate must be specified in the form of a model matrix or formula (respectively). In both cases, we use the same type of formula or model matrix used in the authors’ original publications [18, 19]. For lfdr, the informative covariate must be a discrete group label. We follow the implementation of lfdr developed in [15], which automatically bins the input covariate into 20 approximately equally sized bins before estimating the within-group local FDR (source code to implement this procedure available on the GitHub repository linked in “Data and source code availability”).

Common to all modern FDR-controlling procedures included in Figure 1 is the requirement that the informative covariate also be independent of the *p*-value or test statistic under the null. That is, while the covariate should be informative of whether a test is truly positive, if a test is truly negative, knowledge of the covariate should not alter the validity of the *p*-value or test statistic. Violation of this condition can lead to loss of Type I error control and an inflated rate of false discoveries [30]. To avoid these pitfalls, previously proposed visual diagnostics were used to check both the informativeness and independence of the selected co-variate [15].

### Implementation of benchmarked methods

All analyses were implemented using R version 3.5.0 [24]. We used version 0.99.2 of the R package SummarizedBenchmark [31] to carry out the benchmark comparisons, which is available on GitHub at the ‘fdr-benchmark’ branch at https://github.com/areyesq89/SummarizedBenchmark/tree/fdrbenchmark.

While other modern FDR-controlling methods have also been proposed, methods were excluded if accompanying software was unavailable or if the available software could not be run without substantial work from the user [32].

#### BH

Adjusted *p*-values by BH were obtained using the `p.adjust` function from the `stats` base R package, with option `method=“BH”`.

#### qvalue

Storey’s *q*-values were obtained using the `qvalue` function in version 2.12.0 of the `qvalue` Bioconductor R package.

#### IHW

Adjusted *p*-values by IHW were obtained using the `adj_pvalues` function on the output of the `ihw` function, both from version 1.8.0 of the `IHW` Bioconductor R package.

#### BL

Adjusted *p*-values by BL were obtained by multiplying BH adjusted *p*-values (see above) by the π_0,*i*_ estimates obtained using the `lm_pi0` function from version 1.6.0 of the Bioconductor `R swfdr` package.

#### lfdr

Adjusted *p*-values by lfdr were obtained by first binning the independent covariate into 20 approximately equal sized groups using the `groups_by_filter` function the `IHW R` package. Next the `fdrtool` function from version 1.2.15 of the `fdrtool` CRAN R package was applied to the *p*-values within each covariate bin separately, with the parameter `statistic=“pvalue”`. We require that at least 200 tests per bin, as recommended by `fdrtool`. Note that we follow [15] and use `fdrtool` rather than the `locfdr` package recommended by [17] to obtain local false discovery rates, as the former may operate directly on *p*-values instead of requiring *z*-scores as in the latter.

#### FDRreg

For applications where the test statistic was assumed to be normally distributed, Bayesian FDRs were obtained by `FDRreg` function from version 0.2-1 of the `FDRreg R` package (obtained from GitHub at https://github.com/jgscott/FDRreg). The `features` parameter was specified as a model matrix with a B-spline polynomial spline basis of the independent co-variate with 3 degrees of freedom (using the `bs` function from the `splines` base R package), and no intercept. The `nulltype` was set to `“empirical”` or `“theoretical”` for the empirical and theoretical null implementations of FDRreg, respectively.

#### ASH

*q*-values were obtained using the `get_qvalue` function on the output of the ash function, both from version 2.2-7 of the `ashr R` CRAN package. The effect sizes and their corresponding standard errors were input as the `effect_size` and `sebetahat` parameters, respectively.

#### AdaPT

*q*-values were obtained using the `adapt_glm` function version 1.0.0 of the `adaptMT` CRAN R package. The `pi_formulas` and `mu_formulas` arguments were both specified as natural cubic B-spline basis matrices of the independent covariate with degrees of freedom ∈ {2, 4, 6, 8,10} (using the ns function from the `splines` base R package).

### Yeast *in silico* experiments

Preprocessed RNA-seq count tables from [29] were downloaded from the authors’ GitHub repository at https://github.com/bartongroup/profDGE48. All samples that passed quality control in the original study were included. All genes with a mean count of at least 1 across all samples were included, for a total of 6553 genes. Null comparisons were constructed by randomly sampling two groups of 5 and 10 samples from the same condition (Snf2-knockout). Non-null comparisons of the same size were constructed by adding differentially expressed (DE) genes *in silico* to null comparisons. In addition to different sample sizes, several different settings of the proportion of non-null genes, the distribution of the non-null effect sizes, and informativeness of the covariate were explored. An overview of the different settings is provided in Table S2.

We evaluated results using a low proportion of nonnull genes (500, or approximately 7.5% non-null) as well as a high proportion (2000 or approximately 30% non-null). The non-null genes were selected using probability weights sampled from a logistic function (where weights 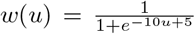, and *u* ~ *U*(0,1)). Three types of informative covariates were explored: (1) strongly informative, (2) weakly informative and (3) uninformative. The strongly informative covariate *X_s_* was equal to the logistic sampling weight *w*. The weakly informative covariate *X_w_* was equal to the logisitic sampling weight plus noise: *w* + *ɛ*, where *ɛ* ~ *N*(0,0.25), truncated such that *X_w_* ∈ (0,1). The uninformative *X_u_* covariate was unrelated to the sampling weights and drawn from a uniform distribution such that *X_u_* ~ *U*(0,1).

We also evaluated results under two different distributions of non-null effect sizes: (1) Unimodal and (2) Bimodal. For unimodal alternative effect size distributions, the observed fold changes for the selected nonnull genes in a non-null empirical comparison of the same sample size were used. For bimodal alternatives, observed test statistics *z* from an empirical non-null comparison of the same sample size were sampled with probability weights *w*(*z*) = *f* (|*x*|; *α*,*β*), where *f* is the Gamma probability density function (with shape and rate parameters *α* = 4.5 and *β* = 1 − 1*e*^−4^, respectively). The corresponding effect sizes (fold changes, *FC*) for ashq were calculated assuming a fixed standard error: *FC* = *zσ_m_*, where *σ_m_* is the median standard error of the *1og*_2_ fold change across all genes.

To add differential signal to the designated nonnull genes, the expression in one randomly selected group was then multiplied by their corresponding fold change. Differential expression analysis using DESeq2 [33] was carried out on both the null and non-null comparisons to assess specificity and sensitivity of the FDR correction methods. Genes for which DESeq2 returned `NA` *p*-values were removed. In each setting, simulations were repeated 100 times and the average and standard error are reported across replications. Results displayed in the main manuscript contain 2000 DE genes, use the strongly informative covariate, and have a sample size of 5 in each group. Results for all settings are presented in Additional files 2-5.

### Polyester *in silico* experiments

The yeast RNA-seq data described in the previous section was used to estimate model parameters using version 1.16.0 of the `polyester` [28] R Bioconductor package. All samples that passed quality control in the original study were included. A baseline group containing all the samples in the wild-type group was used, and genes with mean expression of less than 1 count were filtered out. Counts were library size normalized using DESeq2 size factors [33], and the `get_params` function from the polyester package was used to obtain model parameters. Counts were simulated using the `create_read_numbers` function. Using the same sample sizes as the yeast *in silico* experiments (5 or 10 samples in each group), we evaluated a null comparison, where the `beta` parameters of the `create_read_numbers` function (which represent effect size) were set to zero for all genes. We also evaluated non-null comparisons where the `beta` parameters were drawn from a standard normal distribution for 2000 non-null genes. The non-null genes were selected in the same way as the yeast *in silico* experiments described in the previous section. Differential expression analysis and evaluation of FDR correction methods was also carried out as described for the yeast experiments. Results are presented in Additional file 6.

### Simulation studies

We performed Monte Carlo simulation studies to assess the performance of the methods with known ground truth information. In each simulation, *M* observed effect sizes, 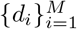, and standard errors, 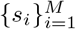, were sampled to obtain test statistics, 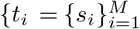. Letting 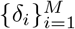 denote the true effect sizes, each *t_i_* was tested against the null hypothesis 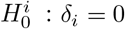. Observed effect sizes and standard errors were simulated to obtain test statistics following one of four distributions under the null:

standard normal distribution,
*t* distribution with 11 degrees of freedom,
*t* distribution with 5 degrees of freedom, or
*χ*^2^ distribution with 4 degrees of freedom.

For each test *i*, let *h_i_* denote the true status of the test, with *h_i_* =0 and *h_i_* = 1 corresponding to the test being null and non-null in the simulation. True effect sizes, *δ_i_*, were set to 0 for {*i*|*h_i_* = 0} and sampled from an underlying non-null effect size distribution for {*i*|*h_i_* = 1}. For normal and *t* distributed test statistics, observed effect sizes were simulated by adding standard normal noise to the true effect sizes, *d_i_ N*(*δ_i_*, 1). The standard errors were all set to 1 to obtain normal test statistics and set to 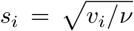 with each 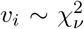 independent to obtain *t* statistics with *v* degrees of freedom. For *χ*^2^ test statistics, observed effect sizes were sampled from non-central *χ*^2^ distributions with non-centrality parameters equal to the true effect sizes, 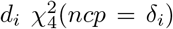. Standard errors were not used to simulate *χ*^2^ statistics and were simply set to 1. The *p*-value was calculated as the two-tail probability of the sampling distribution under the null for normal and *t* statistics. The upper-tail probability under the null was used for *χ*^2^ statistics.

In all simulations, independent covariates, 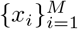, were simulated from the standard uniform distribution over the unit interval. In the uninformative simulation setting, the 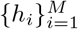 were sampled from a Bernoulli distribution according to the marginal null proportion, 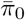, independent of the {*x_i_*}. In all other settings, the 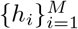 were sampled from Bernoulli distributions with test-specific probabilities determined by the informative covariates through a function, *p*(*x_i_*), taking values in [0,1]. Several forms of *p*(*x_i_*) were considered in the simulations. The *p*(*x_i_*) were chosen to explore a range of relationships between the covariate and the null probability of a test. For further flexibility, the functional relationships were defined conditional on the marginal null probability, 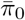, so that similar relationships could be studied across a range of 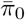. The following 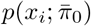 relationships, shown in Additional file 1: Figure S4A for 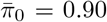, were investigated in the simulations.

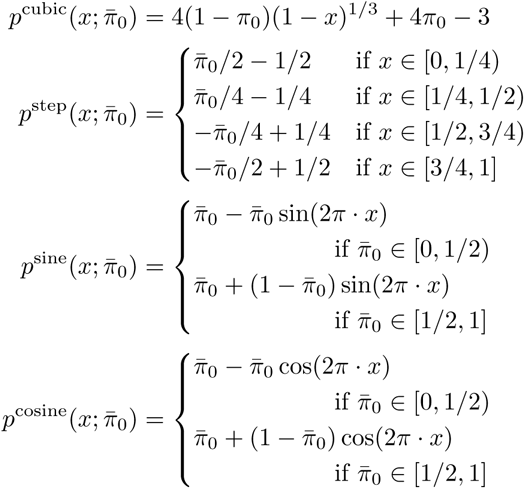

The functions *p*^cubic^ and *p*^step^ are valid, i.e. map to [0,1], for 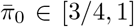 and 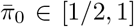, respectively. All others *p*(*x_i_*) are valid for 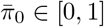. In addition to these functional relationships, we also considered two specialized relationships with 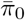 fixed at 0.80. These specialized relationships were parameterized by an informativeness parameter, *δ* ∈ [0,1], such that when *δ* = 0 the covariate was completely uninformative and stratifies the hypotheses more effectively as S increased.

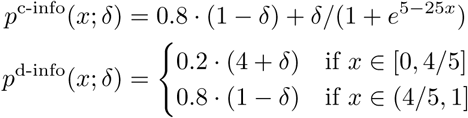

The first, *p*^c-info^, is a continuous relationship between the covariate, *x* and the null proportion, π_0_, while the second, *p*^d-info^, is discrete. Results of simulations with *p*^c-info^ are shown in Additional file 1: Figure S3A-B, where the “informativeness” axis is simply 100 o S. The covariate relationships, *p*^c-info^ and *p*^d-info^ are shown in Additional file 1: Figure S8 across a range of S informativeness values.

Simulations were formulated and performed as several distinct case studies, with full results presented n Additional files 7-20. The complete combination of settings used in each simulation case study are given in Table S3. In each case study, simulations were repeated 100 times, and performance metrics are reported as the average across the replications.

FDRreg and ASH were excluded from simulations with *χ*^2^ distributed test statistics because the setting clearly violated the assumptions of both methods (Figure 1).

### Case studies

We explored several real case-studies using publicly available datasets. Unless otherwise stated, we use an *α* = 0.05 level to define a positive test. In all cases, we also evaluate results using an independent but uninformative covariate (simulated from a standard normal). The locations of where the data can be found and the main details for each case-study is described below. For more specific analysis details, please refer to Additional files 21-41 which contain complete reproducible code sets.

#### Genome-Wide Association Study

GWAS analysis results were downloaded from http://portals.broadinstitute.org/collaboration/giant/images/3/3a/BMI.SNPadjSMK.zip [34]. We used the results subset by European ancestry provided in the file BMI.SNPadjSMK.CombinedSexes.EuropeanO nly.txt to avoid the impact of population stratification on our results. We followed [35] and implemented a linkage disequilibrium (LD)-based pruning step (using the clump command of PLINK v1.90b3s [36]) to remove SNPs in high LD (*r*^2^ < 0.2) with any nearby SNP (<250Kb), based LD estimates from the 1000 Genomes phase three CEU population data [37], available at http://neurogenetics.qimrberghofer.edu.au/iSECA/1000G_20101123_v3_GIANT_chr1_23_minimacnamesifnotRS_CEU_MAF0.01.zip. We explored the use of both sample size and minor allele frequency for each SNP as informative covariates. For fdrreg-e and fdrreg-t, which require a normally distributed test statistic as input, the *t*-statistic (effect size divided by standard error) was used. Because of the large sample sizes in this study (median 161,165), the *t* statistics were approximately normal. For ashq, we used the provided effect size and standard error of the test statistics. Full results are provided in Additional file 21.

#### Gene Set Analysis

We used two RNA-seq datasets that investigated changes in gene expression (1) between cerebellum and cerebral cortex of 5 males from Genotype-Tissue Expression Project [38] and (2) upon differentiation of hematopoietic stem cells (HSCs) into multipotent progenitors (MPP) [39]. For the independent and informative covariate, we considered the size of the gene sets. We considered two different gene set analysis methods: gene set enrichment analysis (GSEA) [40], and overrepresentation testing [41]. To implement the overrepresentation test, we first used version 1.20.0 the `R` Bioconductor package `DESeq2` to obtain a subset of differentially expressed genes (with adjusted *p*-value < 0.05), on which a test of overrepresentation of DE genes among gene sets was performed using version 1.32.0 of the R Bioconductor package `goseq` [41]. To implement GSEA, we used version 1.6.0 of the R Bioconductor package `fgsea` [42], using the DESeq2 test statistics to rank the genes. For both methods, Gene Ontology categories obtained using version 2.36.1 of the R Bioconductor package `biomaRt` containing at least 5 genes were used for the gene sets. For `fgsea`, 10.0 permutations were used and gene sets larger than 500 genes were excluded as recommended in the package documentation. The methods fdrreg-e, fdrreg-t, and ashq were excluded since they require test statistics and/or standard errors that GSEA does not provide. For `goSeq`, we also filtered on gene sets containing at least one DE gene. Full results are provided in Additional file 22-25.

#### Bulk RNA-seq

We used two RNAseq datasets to asses the performance of modern FDR methods in the context of differential expression. The first dataset consisted of 20 paired samples of the *GTEx* project. These 20 samples belonged to two tissues (Nucleus accumbens and Putamen) of 10 female individuals. These samples were preprocessed as described in [43]. We used a second dataset from an experiment in which the microRNA *mir200c* was knocked down in mouse cells [44]. The transcriptomes of knockdown cells and control cells were sequenced in biological duplicates. The processed samples of the knockdown experiment were downloaded from the *recount2* database [45]. For each dataset, we tested for differential expression using `DESeq2`. For FDR methods that can use an informative covariate, we used mean expression across samples, as indicated in the `DESeq2` vignette. Full results are provided in Additional files 26-27.

#### Single-cell RNA-seq

We selected two datasets from the conquer [46] database. First, we detected differentially expressed genes between glioblastoma cells sampled from a tumor core with those from nearby tissue (GSE84465) [47]. In addition, we detected DE genes between murine macrophage cells stimulated to produce an immune response with an unstimulated population (GSE84465) [48]. We filtered out cells with a mapping rate less than 20% or fewer than 5% of genes detected. Genes detected in at least 5% of cells were used in the analysis and spike-in genes were excluded. We carried out DE analyses using two different methods developed for scRNA-seq: scDD [49] and MAST [50], along with the Wilcoxon rank-sum test. MAST was applied to log2(TPM + 1) counts using version 1.6.1 of the `MAST R` Bioconductor package. scDD was applied to raw counts normalized by version 1.8.2 of the `scran` R Bioconductor package [51] using version 1.4.0 of the `scDD` R Bioconductor package. Wilcoxon was applied to counts normalized by TMM (using version 3.22.3 of the `edgeR` R Bioconductor package [52]) using the `wilcox.test` function of the base R package `stats`. We examined the mean nonzero expression and detection rate (defined as the proportion of cells expressing a given gene) as potentially informative covariates. The fdrreg-e and fdrreg-t methods were excluded since none of the test statistics used are normally distributed. Likewise, ashq was excluded since none of the methods considered provide effect sizes and standard errors. Full results are provided in Additional files 28-33.

#### ChIP-seq

ChIP-seq analyses were carried out on two separate datasets. First, H3K4me3 data from two cell lines (GM12878 and K562) were downloaded from ENCODE portal [53]. In each cell line, four replicates were selected with half of them from one laboratory and the other half from another laboratory. We performed two types of differential binding analyses between cell lines: (1) Following the workflow of [54], we used the csaw [55] method (version 1.14.1 of the `csaw` Bioconductor R package) to identify candidate *de novo* differential windows, and edgeR [52] method (using version 3.22.3 of the `edgeR` R Bioconductor package) to test for significance. (2) We tested for differential binding only in predefined promoter regions (UCSC “Known Gene” annotations for human genome assembly hg19 (GRCh37) using DESeq2 [33]). In addition, we carried out analysis type (1) on a second ChIP-seq dataset to compare CREB-binding protein in wild-type and CREB knockout mice [54, 56] (GSE54453). For analysis type (1), we used the region width as the informative covariate. For analysis type (2), we used mean coverage as the informative covariate. The fdrreg-e and fdrreg-t methods were excluded from csaw analyses since the test statistics used are non normally distributed. Likewise, ashq was excluded since csaw does not provide effect sizes and standard errors. Full results are provided in Additional files 34-36.

#### Microbiome

We performed two types of analyses (1) differential abundance analysis, and (2) correlation analysis. For the differential abundance analyses, we used four different datasets from the MicrobiomeHD database [57]: (1) obesity [58], (2) inflammatory bowel disease (IBD) [59], (3) infectious diarrhea (Clostridium difficile (CDI) and non-CDI) [60], and (4) colorectal cancer (CRC) [61]. These studies were processed as described in [57].

We performed Wilcoxon rank-sum differential abundance tests on the operational taxonomic units (OTUs, sequences clustered at 100% genetic similarity) and on taxa collapsed to the genus level (as in [57]). Full results are provided in Additional files 37-40.

For the correlation analyses, we used a previously published dataset of microbial samples from monitoring wells in a site contaminated by former waste disposal ponds, where all sampled wells have various geochemical and physical measurements [62]. Paired-end reads were merged using PEAR (version 0.9.10) and demultiplexed with QIIME v 1.9.1 split_libraries_fastq.py (maximum barcode error of 0 and quality score cutoff of 20) [63, 64]. Reads were dereplicated using USEARCH v 9.2.64 -fastx_uniques and operational taxonomic units (OTUs) were called with -cluster_otus and an identity threshold of 0.97 [65]. These data were processed with the Amplicon Sequence Analysis Pipeline http://zhoulab5.rccc.ou.edu/pipelines/ASAP_web/pipeline_asap.php. We carried out a Spearman correlation test (*H*_0_ : *ρ* = 0) between OTU relative abundances across wells and the respective values of three geochemical variables: pH, Al, and SO_4_. Full results are provided in Additional file 41.

For all analyses we examined the ubiquity (defined as the percent of samples with non-zero abundance of each taxa) and mean non-zero abundance of taxa as potentially informative covariates. Results in the main manuscript are shown for the OTU level, unless there were no rejections in the vast majority of methods, and then results are shown for genus level. The SO_4_ dataset was also excluded from the main results since most methods find no rejections. We excluded fdrreg-e, fdrreg-t, and ashq since neither the Wilcoxon nor Spearman test statistics are normally distributed, nor do they provide an effect size and standard error. Due to small numbers of tests, lfdr was excluded from the obesity and IBD genus level analyses.

### Evaluation metrics

All studies (*In-silico* experiments, simulations, and case studies) were evaluated on the number of rejections at varying *α* levels, ranging from 0.01 to 0.10. The overlap among rejection sets for each method was examined using version 1.3.3 of the `UpSetR` R CRAN package. *In-silico* experiments and simulation studies were also evaluated on the following metrics at varying *α* levels, ranging from 0.01 to 0.10: True Positive Rate (TPR), observed FDR, and True Negative Rate (TNR). Here we define TPR as the number of true positives out of the total number of non-null tests, observed FDR as the number of false discoveries out of the total number of discoveries (defined as 0 when there are no discoveries), and TNR as the number of true negatives out of the total number of null tests.

### Summary metrics

The final ratings presented in Figure 6 were determined from the aggregated results of the simulations, yeast experiments, and case studies.

#### FDR control

Ratings were determined using results from non-null simulations where all methods were applied (i.e. excluding *χ*^2^ settings), and all non-null yeast experiments. In each setting or experiment, a method was determined to control FDR at the nominal 5% cutoff if the mean FDR across replications was less than one standard error above 5%. The following cutoffs were used to determine superior, satisfactory, and unsatisfactory methods.

Superior: failed to control FDR in less than 10% of settings in both simulations and yeast experiments.
Satisfactory: failed to control FDR in less than 10% of settings in either simulations or yeast experiments.
unsatisfactory: otherwise.

The computed proportion of simulation and yeast settings exceeding the nominal FDR threshold are shown in Figure 5A.

#### Power

Similar to above, ratings were determined using results from non-null simulations where all methods were applied (i.e. excluding *χ*^2^ settings), and all non-null yeast experiments. In each setting or experiment, methods were ranked in descending order according to the mean TPR across replications at the nominal 5% FDR cutoff. Ties were set to the intermediate value, e.g 1.5, if two methods tied for the highest TPR. The mean rank of each method was computed across simulation settings and yeast experiments separately, and used to determine superior, satisfactory, and unsatisfactory methods according to the following cutoffs.

Superior: mean TPR rank less than 5 (of 8) in both simulations and yeast experiments.
Satisfactory: mean TPR rank less than 6 (of 8) in both simulations and yeast experiments.
Unsatisfactory: otherwise.

Mean TPR ranks for methods across simulation and yeast settings are shown in Figure 5B.

#### Consistency

Ratings were determined using results from non-null simulations where all methods were applied (i.e. excluding *χ*^2^ settings) and all case studies. Here, ashq and fdrreg-t were excluded from metrics computed using the case studies, as the two methods were only applied in 4 of the 26 case studies. In each simulation setting, the TPR and FDR of each covariate-aware method was compared against the TPR and FDR of both the BH approach and Storey’s *q*-value at the nominal 5% FDR cutoff. Similarly, in each case study, the number of rejections of each method was compared against the number of rejections of BH and Storey’s *q*-value. Based on these comparisons, two metrics were computed for each method.

First, in each setting and case study, a modern method was determined to underperform the classical approaches if the TPR or number of rejections was less than 95% of the minimum of the BH approach and Storey’s *q*-value. The proportion of simulation settings and case studies where a modern method underperformed was used to determine the consistent stability of an approach (Figure 5C). The FDR of the methods was not used for this metric.

Second, in each simulation setting and case study, the log-ratio FDR, TPR and number of rejections was computed against a baseline for each modern method. For each setting, the average across the classical methods was used as the baseline. Methods were then ranked according to the standard deviation of these log-ratios, capturing the consistency of the methods across simulations and case studies (Figure 5D).

These two metrics were used to determine superior, satisfactory, and unsatisfactory methods according to the following cutoffs.

Superior: In top 50% (3) of methods according to variance of log-ratio metrics.
Satisfactory: Not in top 50% (3) of methods according to variance of log-ratio metrics, but underperformed both BH and Storey’s *q*-value in less than 10% of case studies or simulation settings.
Unsatisfactory: Not in top 50% (3) of methods according to variance of log-ratio metrics, and underperformed both BH and Storey’s *q*-value in more than 10% of case studies or simulation settings.

#### Applicability

Ratings were determined using the proportion of case studies in which each method could be applied. The proportion was calculated first within each type of case studies, followed by averaging across all case studies. This was done to prevent certain types, e.g. scRNA-seq DE analysis, from dominating the average. The following cutoffs were used to determine superior, satisfactory, and unsatisfactory methods. For this metric, cases when adapt-glm returned exactly zero positive tests while all other methods returned non-zero results where labeled as data sets where the method could not be applied. This is denoted by an asterisk in Figure 6.

Superior: applied in 100% of case studies.
Satisfactory: applied in more than 50% of case studies.
Unsatisfactory: otherwise.

#### Usability

Ratings were determined based on our experience using the method for our benchmark comparison, and rated according to the following criteria.

Superior: a well-documented implementation is available.
Satisfactory: an implementation is available, but lacks extensive documentation or requires additional work to install.
Unsatisfactory: no implementation is readily available.

### Data and source code availability

Full analyses and benchmarking methods of the *in silico* experiments, simulations, and case studies are provided in Additional files 2-41. The source code to reproduce all results in the manuscript and additional files, as well as all figures is available on GitHub (https://github.com/pkimes/benchmark-fdr). The SummarizedBenchmark package is available on Bioconductor. The Ecosystems and Networks Integrated with Genes and Molecular Assemblies (ENIGMA) data used in the microbiome case study is available at https://zenodo.org/record/1455793/. For the case-studies, the data sources are described in detail in the Methods section.

## Competing interests

The authors declare they have no competing interests.

## Author’s contributions

All authors conceptualized and contributed to the main aims of the project. PKK, SCH, and KK performed the simulation studies. For the case-studies: CD performed the testing in microbiome data, KK performed the GWAS data analysis, AR and KK performed the gene set analyses, CS, PKK, and AR performed the bulk RNA-seq data analysis, AS and KK performed the single-cell RNA-seq data analysis, MT and PKK performed the ChIP-seq data analysis. All authors wrote and approved the final manuscript.

## Acknowledgements

We are grateful to Rafael A. Irizarry for valuable suggestions on method evaluation and results presentation. We also thank Nikolaos Ignatiadis, James Scott, Lihua Lei, Mengyin Lu, Peter Carbonetto, Matthew Stephens, and Simina Boca for useful suggestions and help with inquiries about the usage of specific methods. Finally, we also thank Michael I. Love for suggestions on modern FDR methods to incorporate into our study.

## Tables

### Additional files

Additional file 1 — **Supplementary Figures, Tables, and Results.** Supplementary Figures S1-S13, Supplementary Tables S1-S4, and Supplementary results (PDF).

Additional file 2 — **Yeast *in silico* experiments I.** Analysis and benchmarking results under the null, and using a unimodal alternative effect size distribution and large proportion (30%) of non-nulls using yeast RNA-seq data (HTML).

Additional file 3 — **Yeast *in silico* experiments II.** Analysis and benchmarking results using a unimodal alternative effect size distribution and small proportion (7.5%) of non-nulls using yeast RNA-seq data (HTML).

Additional file 4 — **Yeast *in silico* experiments III.** Analysis and benchmarking results using a bimodal alternative effect size distribution and large proportion (30%) of non-nulls using yeast RNA-seq data (HTML).

Additional file 5 — **Yeast *in silico* experiments IV.** Analysis and benchmarking results using a bimodal alternative effect size distribution and small proportion (7.5%) of non-nulls using yeast RNA-seq data (HTML).

Additional file 6 — **Polyester *in silico* experiments.** Analysis and benchmarking results using a bimodal alternative effect size distribution and large proportion (30%) of non-nulls using RNA-seq data simulated using Polyester (HTML).

Additional file 7 — **Simulations I: Null.** Analysis and benchmarking results of simulation settings with only null tests, using normal, *t*, and *χ*^2^ distributed test statistics (HTML).

Additional file 8 — **Simulations II: Informative (cubic).** Analysis and benchmarking results of simulation settings with “cubic” informative covariate and normal, *t*, and *χ*^2^ distributed test statistics (HTML).

Additional file 9 — **Simulations III: Informative (step).** Analysis and benchmarking results of simulation settings with “step” informative covariate and normal, *t*, and *χ*^2^ distributed test statistics (HTML).

Additional file 10 — **Simulations IV: Informative (sine).** Analysis and benchmarking results of simulation settings with “sine” informative covariate and normal, *t*, and *χ*^2^ distributed test statistics (HTML).

Additional file 11 — **Simulations V: Informative (cosine).** Analysis and benchmarking results of simulation settings with “cosine” informative covariate and normal, *t*, and *χ*^2^ distributed test statistics (HTML).

Additional file 12 — **Simulations VI: Unimodal Effect Sizes.** Analysis and benchmarking results of simulation settings with “cubic” informative covariate, normal test statistics and unimodal effect size distributions (HTML).

Additional file 13 — **Simulations VII: Unimodal Effect Sizes** (*t*_11_ **test statistics).** Analysis and benchmarking results of simulation settings with “cubic” informative covariate, *t*_11_ distributed test statistics and unimodal effect size distributions (HTML).

Additional file 14 — **Simulations VIII: Unimodal Effect Sizes (25% non-null).** Analysis and benchmarking results of simulation settings with “cubic” informative covariate, normal test statistics, unimodal effect size distributions, and higher (25% vs 10%) proportion of non-null tests (HTML).

Additional file 15 — **Simulations IX: Varying *M* Tests.** Analysis and benchmarking results of simulation settings with “sine” informative covariate, normal test statistics, and varying total number of hypothesis tests (HTML).

Additional file 16 — **Simulations X: Varying Null Proportion.** Analysis and benchmarking results of simulation settings with “sine” informative covariate, normal test statistics, and varying proportion of null hypotheses (HTML).

Additional file 17 — **Simulations XI: Varying Null Proportion** (*t*_11_ **test statistics).** Analysis and benchmarking results of simulation settings with “sine” informative covariate, *t*_11_ distributed test statistics, and varying proportion of null hypotheses (HTML).

Additional file 18 — **Simulations XII: Varying Informativeness (continuous** *p*(*x*; (*δ*)). Analysis and benchmarking results of simulation settings with informative covariates of varying informativeness using a continuous relationship between the covariate and the null proportion (HTML).

Additional file 19 — **Simulations XIII: Varying Informativeness (discrete** *p*(*x*; (*δ*)). Analysis and benchmarking results of simulation settings with informative covariates of varying informativeness using a discrete relationship between the covariate and the null proportion (HTML).

Additional file 20 — **Simulations XIV: AdaPT with null option.** Analysis and benchmarking results of simulation settings with “step” informative covariate comparing AdaPT with and without a null model option (HTML).

Additional file 21 — **Case study: Genome-wide association studies.** Analysis and benchmarking results of a meta-analysis of Genome-wide association studies of body mass index (HTML).

Additional file 22 — **Case study: Gene set analyses I.** Gene set analysis of differentially expressed mouse genes after HSC differentiation using goSeq (HTML).

Additional file 23 — **Case study: Gene set analyses II.** Gene set analysis of differentially expressed human genes between cerebellum and cerebral cortex using goSeq (HTML).

Additional file 24 — **Case study: Gene set analyses III.** Gene set analysis and benchmarking results of differentially expressed mouse genes after HSC differentiation using GSEA (HTML).

Additional file 25 — **Case study: Gene set analyses IV.** Gene set analysis and benchmarking results of differentially expressed human genes between cerebellum and cerebral cortex using GSEA (HTML).

Additional file 26 — **Case study: Differential gene expression in bulk RNA-seq I.** Differential expression analysis and benchmarking results of human genes between Nucleus accumbens and Putamen (HTML).

Additional file 27 — **Case study: Differential gene expression in bulk RNA-seq II.** Differential expression analysis and benchmarking results of human genes between microRNA knockdown and control mouse cells (HTML).

Additional file 28 — **Case study: Differential gene expression in single-cell RNA-seq I.** Single-cell differential expression analysis and benchmarking results between human glioblastoma tumor cells and nearby controls using MAST (HTML).

Additional file 29 — **Case study: Differential gene expression in single-cell RNA-seq II.** Single-cell differential expression analysis and benchmarking results between human glioblastoma tumor cells and nearby controls using scDD (HTML).

Additional file 30 — **Case study: Differential gene expression in single-cell RNA-seq III.** Single-cell differential expression analysis and benchmarking results between human glioblastoma tumor cells and nearby controls using Wilcox (HTML).

Additional file 31 — **Case study: Differential gene expression in single-cell RNA-seq IV.** Single-cell differential expression analysis and benchmarking results between stimulated murine macrophage cells and controls using MAST (HTML).

Additional file 32 — **Case study: Differential gene expression in single-cell RNA-seq V.** Single-cell differential expression analysis and benchmarking results between stimulated murine macrophage cells and controls using scDD (HTML).

Additional file 33 — **Case study: Differential gene expression in single-cell RNA-seq VI.** Single-cell differential expression analysis and benchmarking results between stimulated murine macrophage cells and controls using Wilcox (HTML).

Additional file 34 — **Case study: Differential binding in ChIP-seq I.** Differential binding analysis and benchmarking results of H3K4me3 between two cell lines in promoter regions using edgeR (HTML).

Additional file 35 — **Case study: Differential binding in ChIP-seq II.** Differential binding analysis and benchmarking results of H3K4me3 between two cell lines using csaw (HTML).

Additional file 36 — **Case study: Differential binding in ChIP-seq III.** Differential binding analysis and benchmarking results of CREB-binding protein between knockout and wild-type mice using csaw (HTML).

Additional file 37 — **Case study: Differential abundance testing in microbiome data analysis I.** Differential abundance analysis and benchmarking results of obesity (HTML).

Additional file 38 — **Case study: Differential abundance testing in microbiome data analysis II.** Differential abundance analysis and benchmarking results of IBD (HTML).

Additional file 39 — **Case study: Differential abundance testing in microbiome data analysis III.** Differential abundance analysis and benchmarking results of infectious diarrhea (HTML).

Additional file 40 — **Case study: Differential abundance testing in microbiome data analysis IV.** Differential abundance analysis and benchmarking results of colorectal cancer (HTML).

Additional file 41 — **Case study: Correlation testing in microbiome data analysis.** Correlation analysis and benchmarking results of wastewater contaminants (HTML).

### Additional file 1

#### Supplementary Figures and Tables

**Figure S1.**
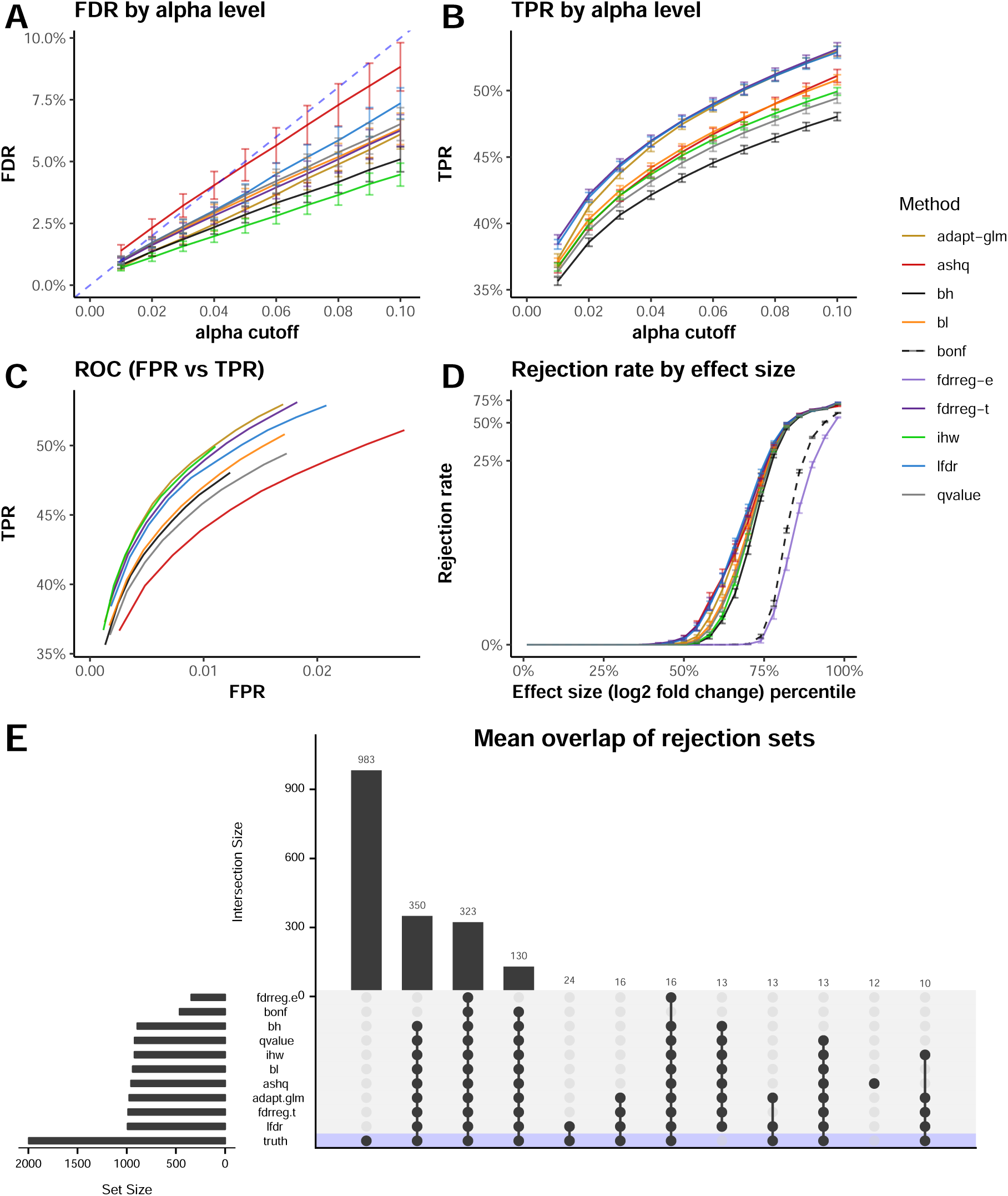
Performance evaluation in yeast *in silico* experiment. Using RNA-seq data derived from a yeast experiment, a differential expression analysis was performed comparing two groups with five samples each and differential signal added to 2000 out of approximately 6500 genes (30%) using an informative covariate. An evaluation of the performance includes the (A) FDR, (B) TPR, and (C) number of rejections by FDR (alpha) cutoff. (D) Percentage significant at 0.05 cutoff (log-scale) by effect size (log2 fold change) percentile. Method is denoted by color and the mean value across 100 replications with standard error bars is shown in (A)-(D). (E) Mean overlap of discoveries at 0.05 cutoff across methods and underlying truth.

**Figure S2.**
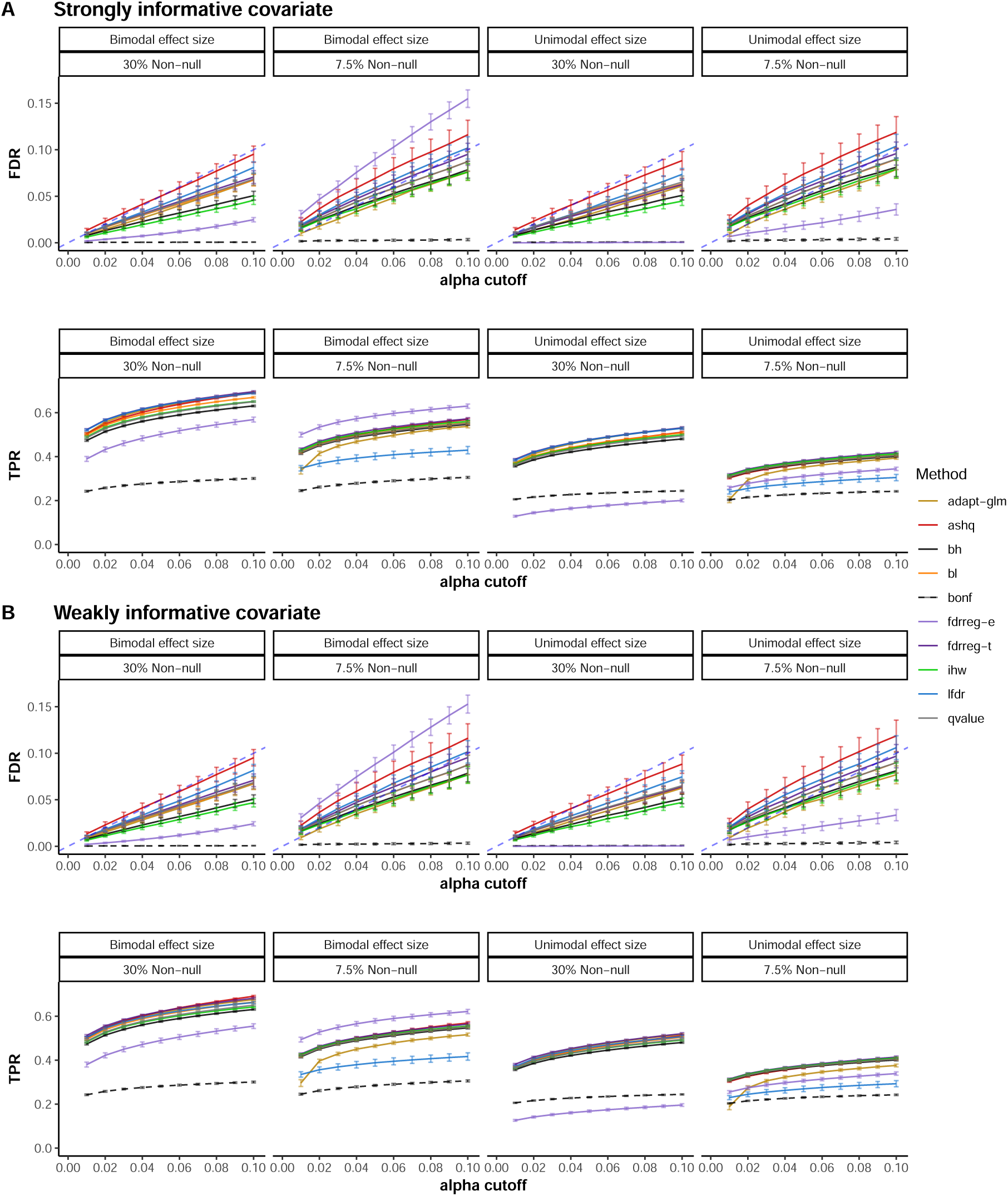
FDR and TPR under various spike-in settings of yeast *in silico* experiments. Plots of FDR and TPR across *α* cutoff values over 100 replications in the yeast simulation (sample size 5 in each group) by alpha level. Vertical bars depict standard error. Each panel within A and B represents a combination of settings for π_0_: 30% (2000) and 7.5% (500) non-null genes (total of 6500 genes), as well as different non-null effect size distributions: bimodal and unimodal. (A) For a *strongly informative covariate*: the informative covariate is equal to the sampling weights for non-null genes. (B) For a *weakly informative covariate*: the informative covariate is equal to the sampling weights for selecting non-null genes plus noise.

**Figure S3.**
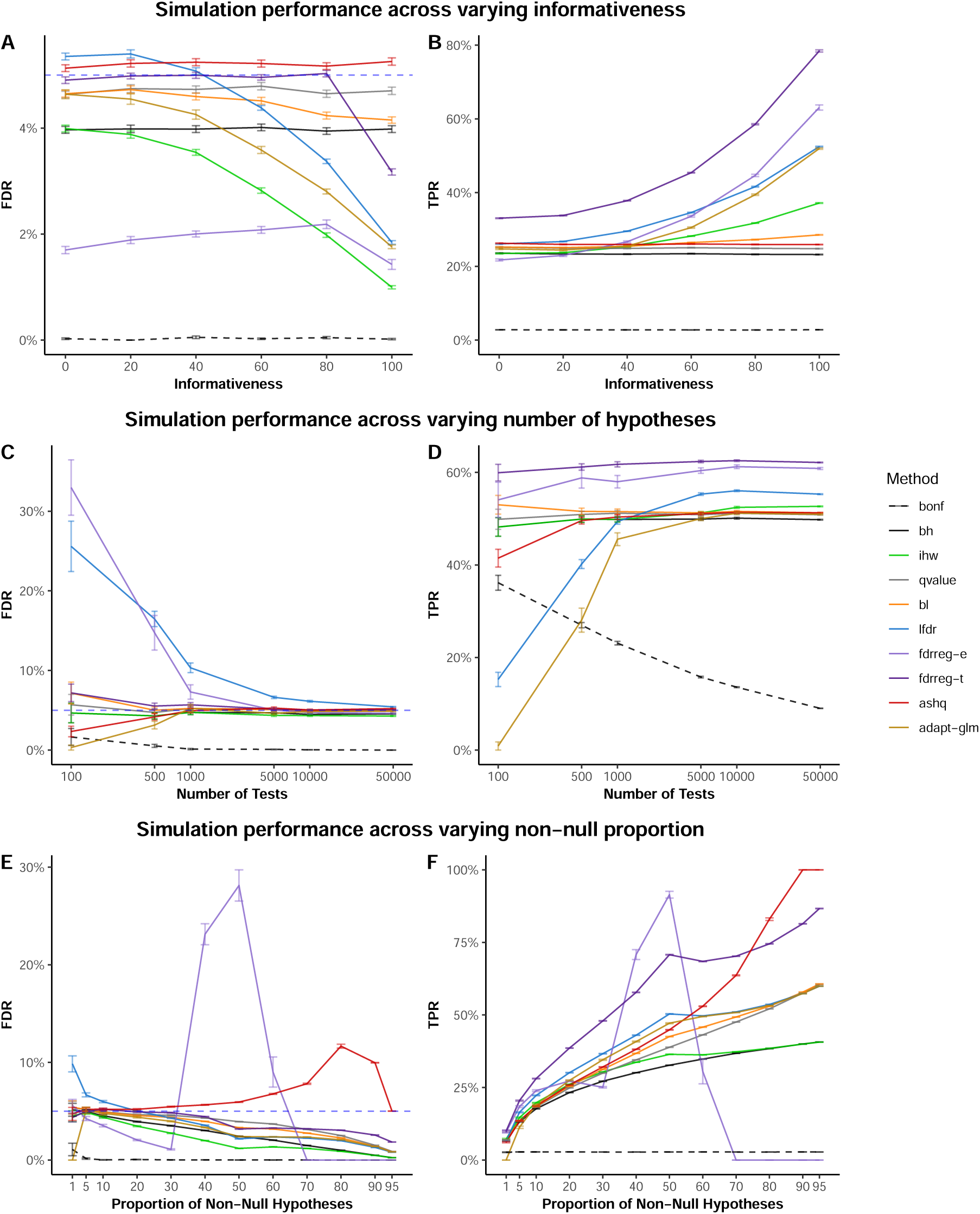
Performance evaluation in simulations. (A) FDR control and (B) TPR across varying informativeness. (C) FDR and (D) TPR across varying number of hypotheses (E) FDR and (F) TPR across varying proportions of null and alternative hypotheses. Method is denoted by color and the mean value across 100 replications with standard error bars is shown in (A)-(F). The FDR and TPR are shown at the nominal 0.05 cutoff.

**Figure S4.**
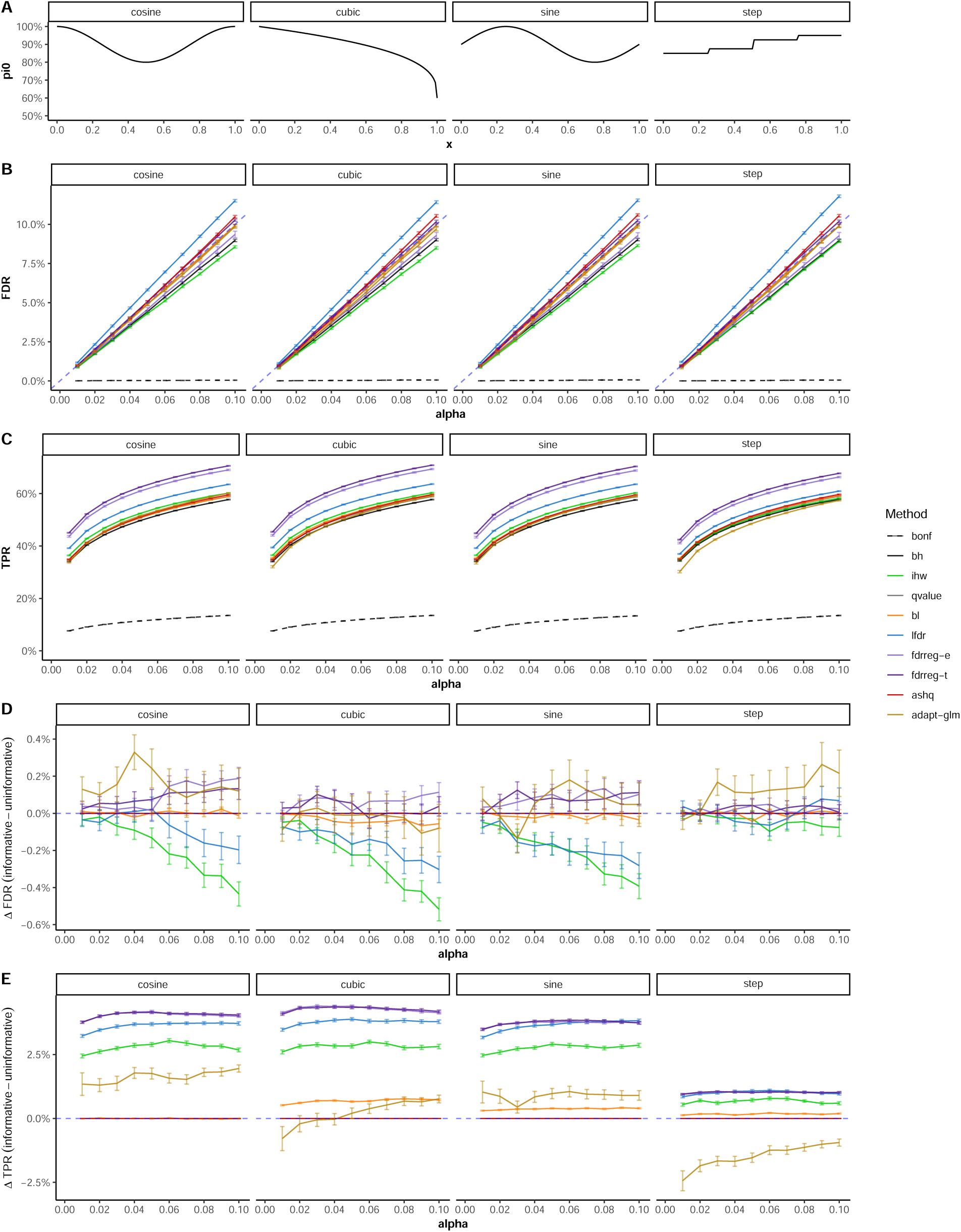
Simulation performance across informative covariate relationship. (A) Relationship between covariate value, x, and null proportion of hypotheses, π_0_, across simulation settings. (B) FDR and (C) TPR across nominal FDR thresholds between 0.01 and 0.10 are shown for each method across four informative covariates described in the “Methods” section. Differences in (D) FDR and (E) TPR between informative and uninformative covariates are also shown.

**Figure S5.**
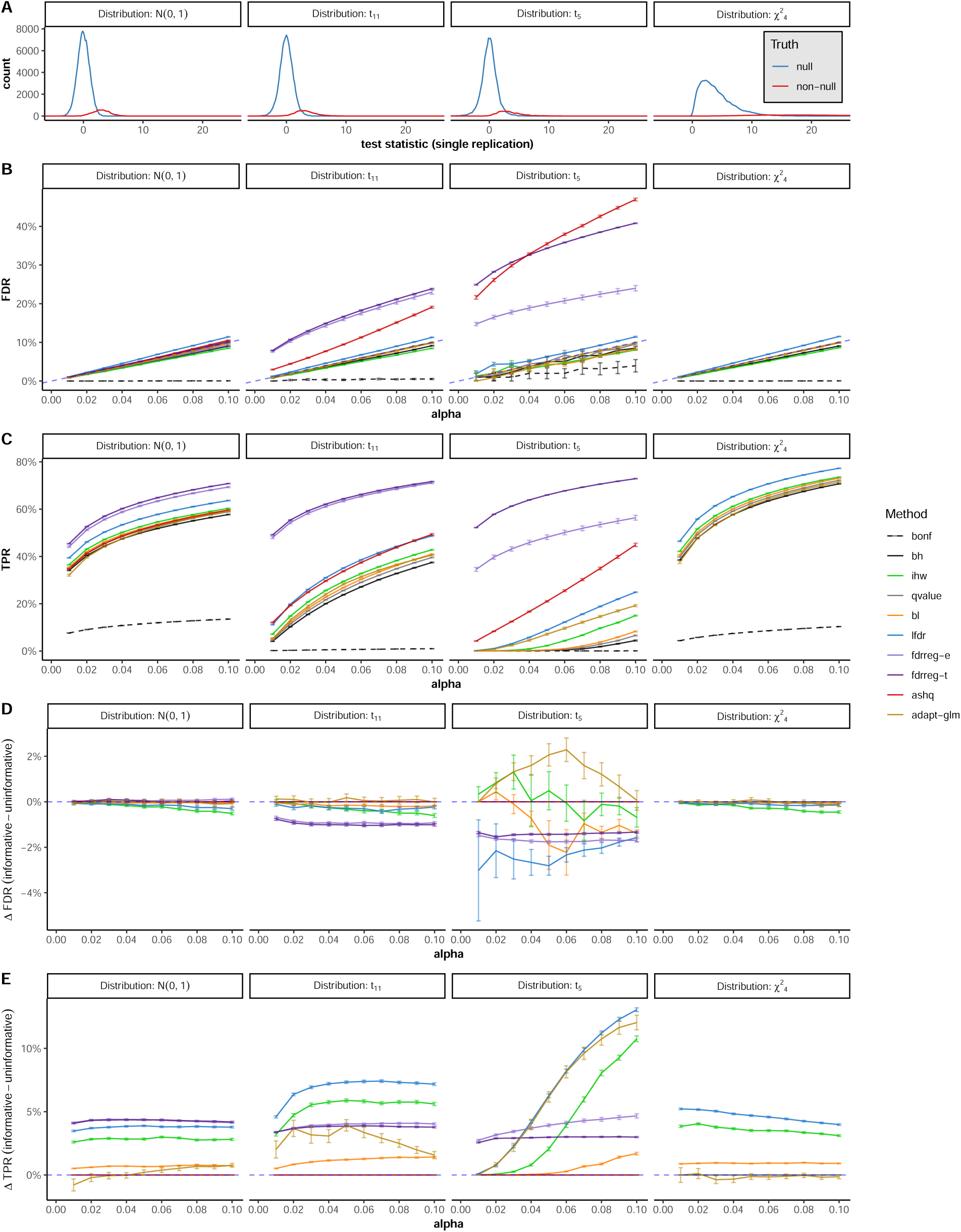
Simulation performance across test statistic distributions. (A) Distributions of test statistics for null and non-null tests in one replication of each simulation. (B) FDR control and (C) TPR across different test statistic distributions. Differences in (D) FDR and (E) TPR between informative and uninformative covariates are also shown. Method is denoted by color and the mean value across 100 replications with standard error bars is shown in (B)-(E). Results for ash, fdrreg-t, fdrreg-e are not shown for the 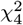 setting due to due to distributional assumptions of the methods (see Table 1). Method is denoted by color and the mean value across 100 replications with standard error bars is shown in (B)-(E).

**Figure S6.**
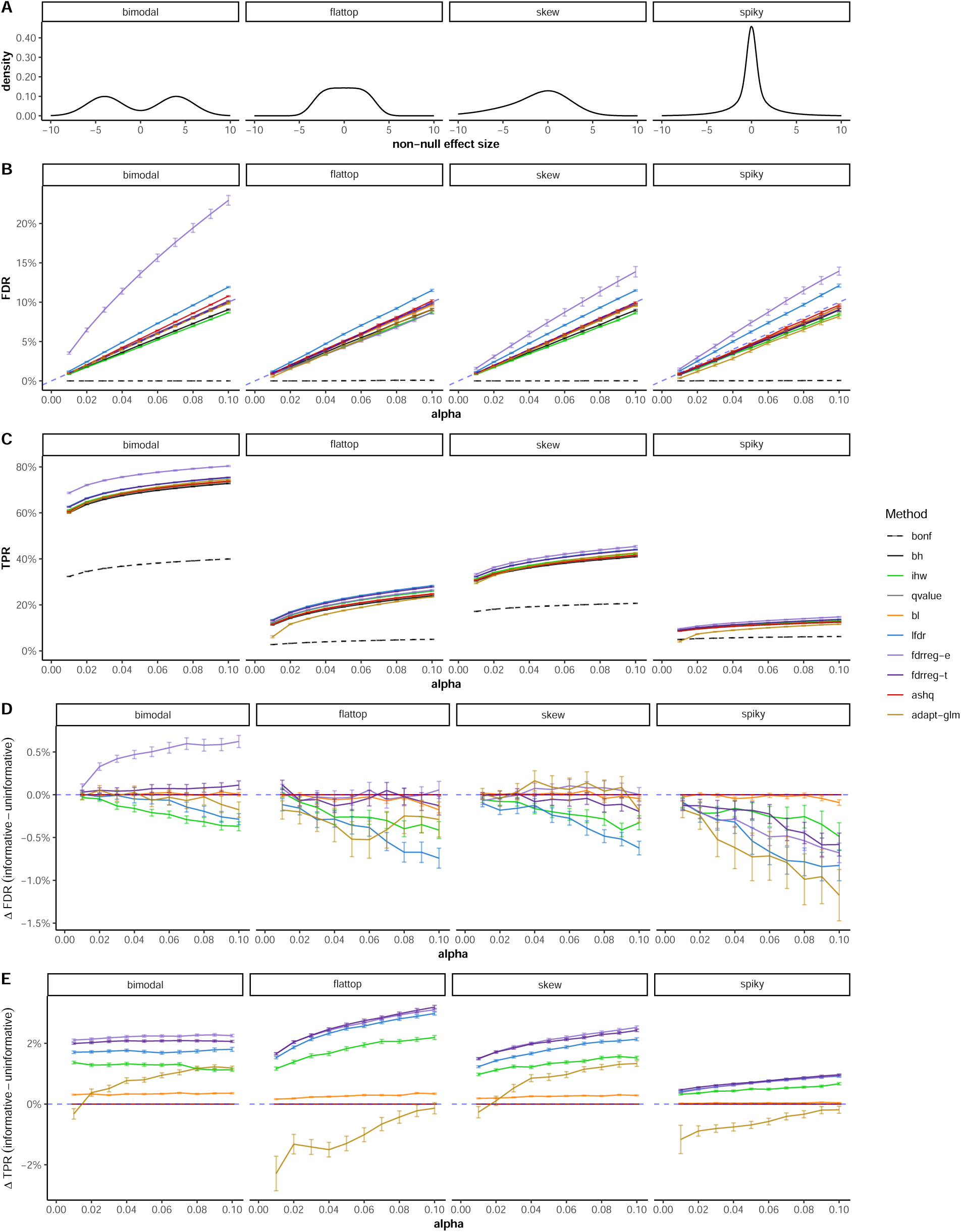
Simulation performance across effect size distributions. (A) Distributions of effect sizes included in unimodal effect size simulations. (B) FDR and (C) TPR across nominal FDR thresholds between 0.01 and 0.10 are shown for each method across four distributions of the non-null effect sizes presented in [20]: bimodal, flat-top, skew, and spiky. All distributions are unimodal with mode at zero except for the “bimodal” setting. Settings are tested to evaluate the performance of ASH against all other methods under the unimodal assumption. Differences in (D) FDR and (E) TPR between informative and uninformative covariates are also shown. Method is denoted by color and the mean value across 100 replications with standard error bars is shown in (B)-(E).

**Figure S7.**
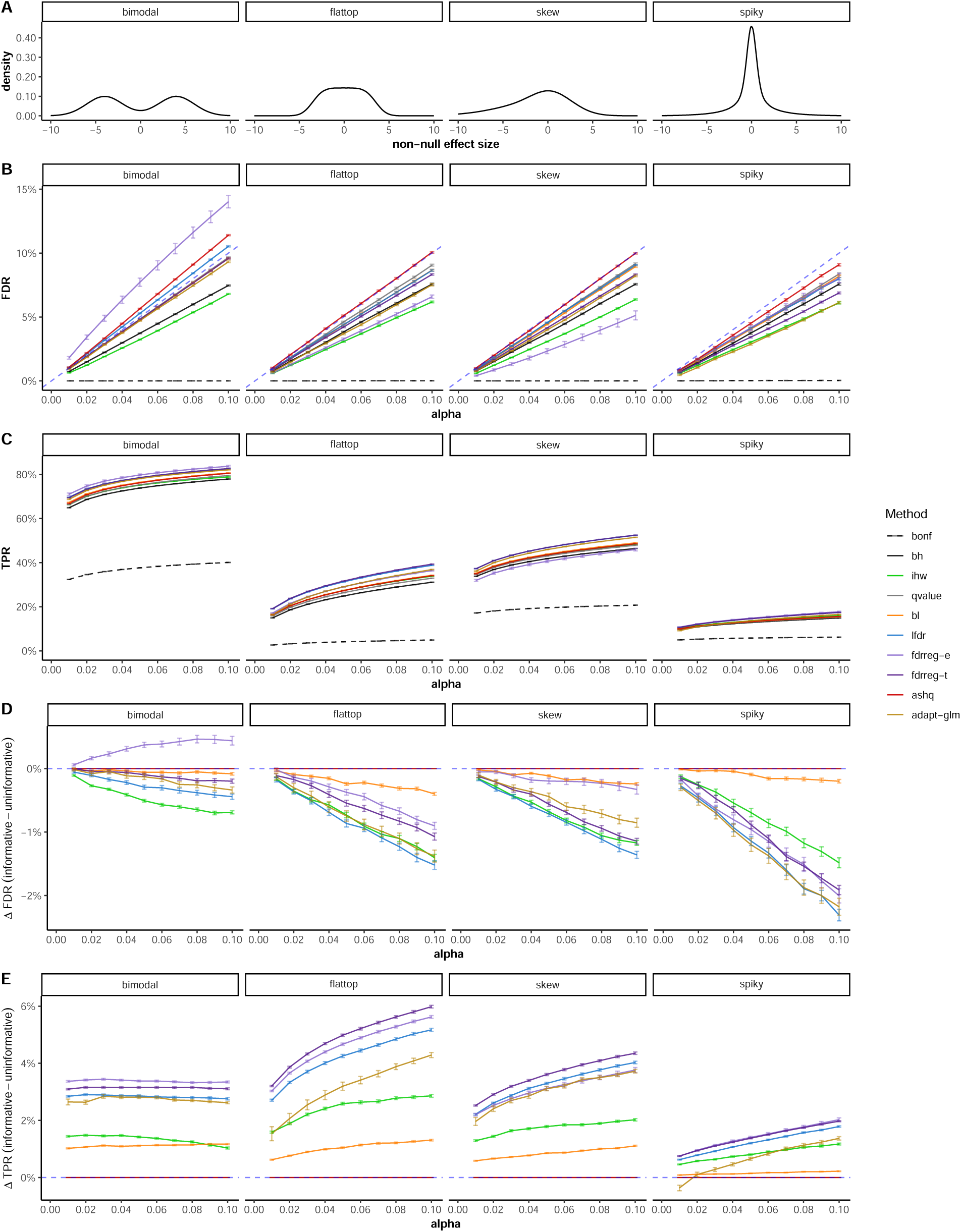
Simulation performance across unimodal effect size distributions w/ 25% non-null. Same as Figure S6 but with increased proportion of non-null hypotheses. (A) Distributions of effect sizes included in unimodal effect size simulations. (B) FDR and (C) TPR across nominal FDR thresholds between 0.01 and 0.10 are shown for each method across four distributions of the non-null effect sizes. Differences in (D) FDR and (E) TPR between informative and uninformative covariates are also shown. Method is denoted by color and the mean value across 100 replications with standard error bars is shown in (B)-(E).

**Figure S8.**
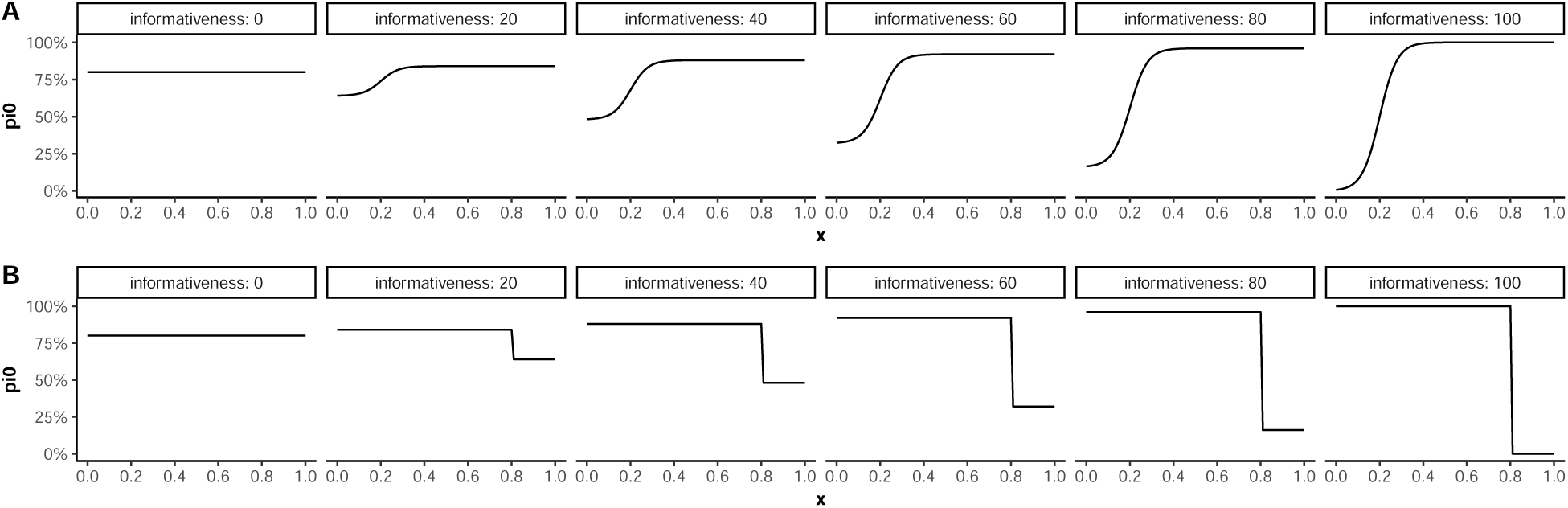
Informativeness covariate relationships. Relationship between covariate value, x, and null proportion of hypotheses, π_0_, across several *δ* informativeness values for (A) *p*^c-info^ and (B) *p*^s-info^.

**Figure S9.**
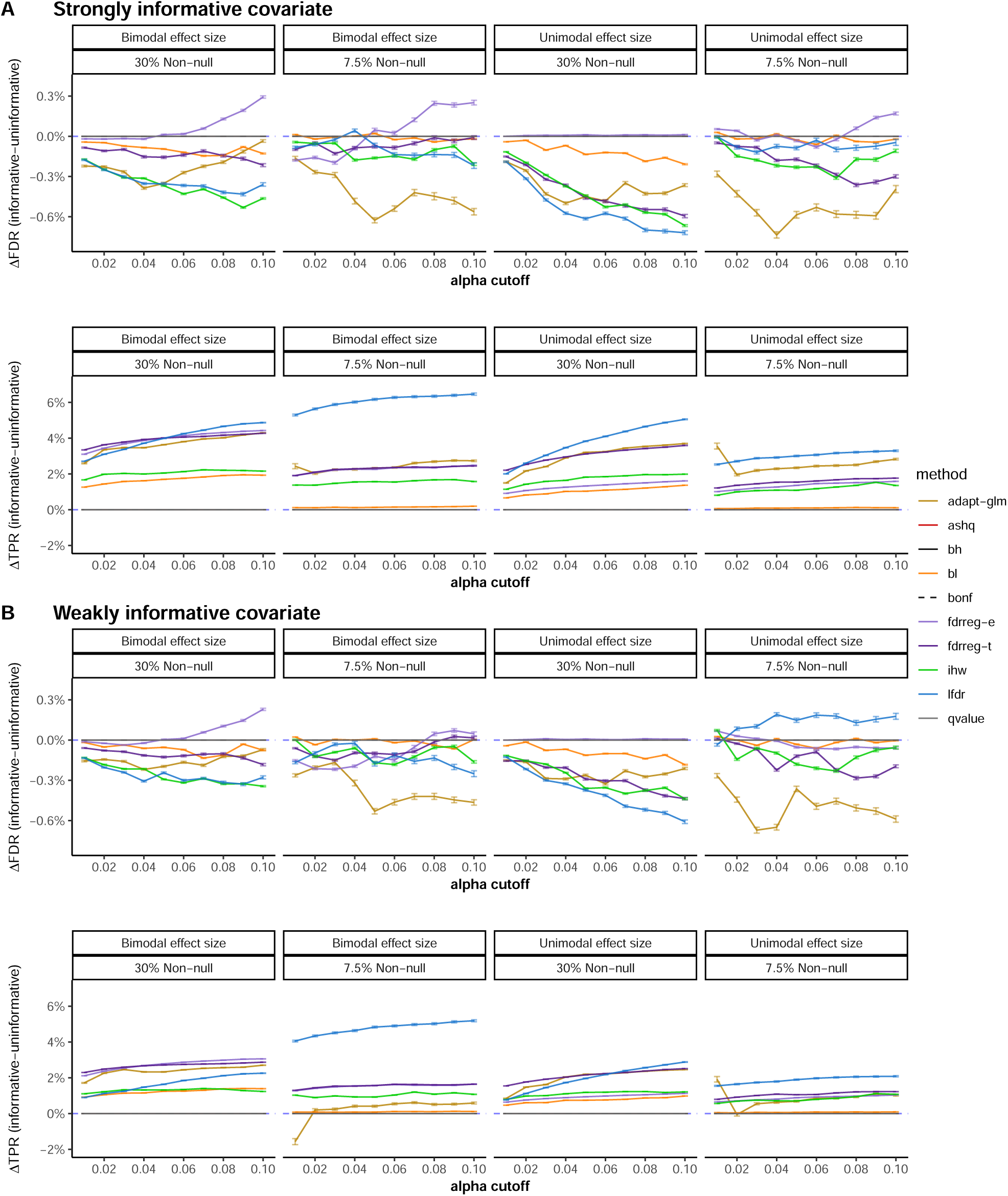
Impact of covariate information in yeast experiments. Mean difference of FDR and TPR (with informative covariate - without informative covariate) over 100 replications in the yeast simulation (sample size 5 in each group) by alpha level. Vertical bars depict standard error. Each panel within A and B represents a combination of settings for π_0_: 30% (2000) and 7.5% (500) non-null genes (total of 6500 genes), as well as different non-null effect size distributions: bimodal and unimodal. (A) For a *strongly informative covariate*: the informative covariate is equal to the sampling weights for non-null genes, and a (B) For a *weakly informative covariate*: the informative covariate is equal to the sampling weights for selecting non-null genes plus noise.

**Figure S10.**
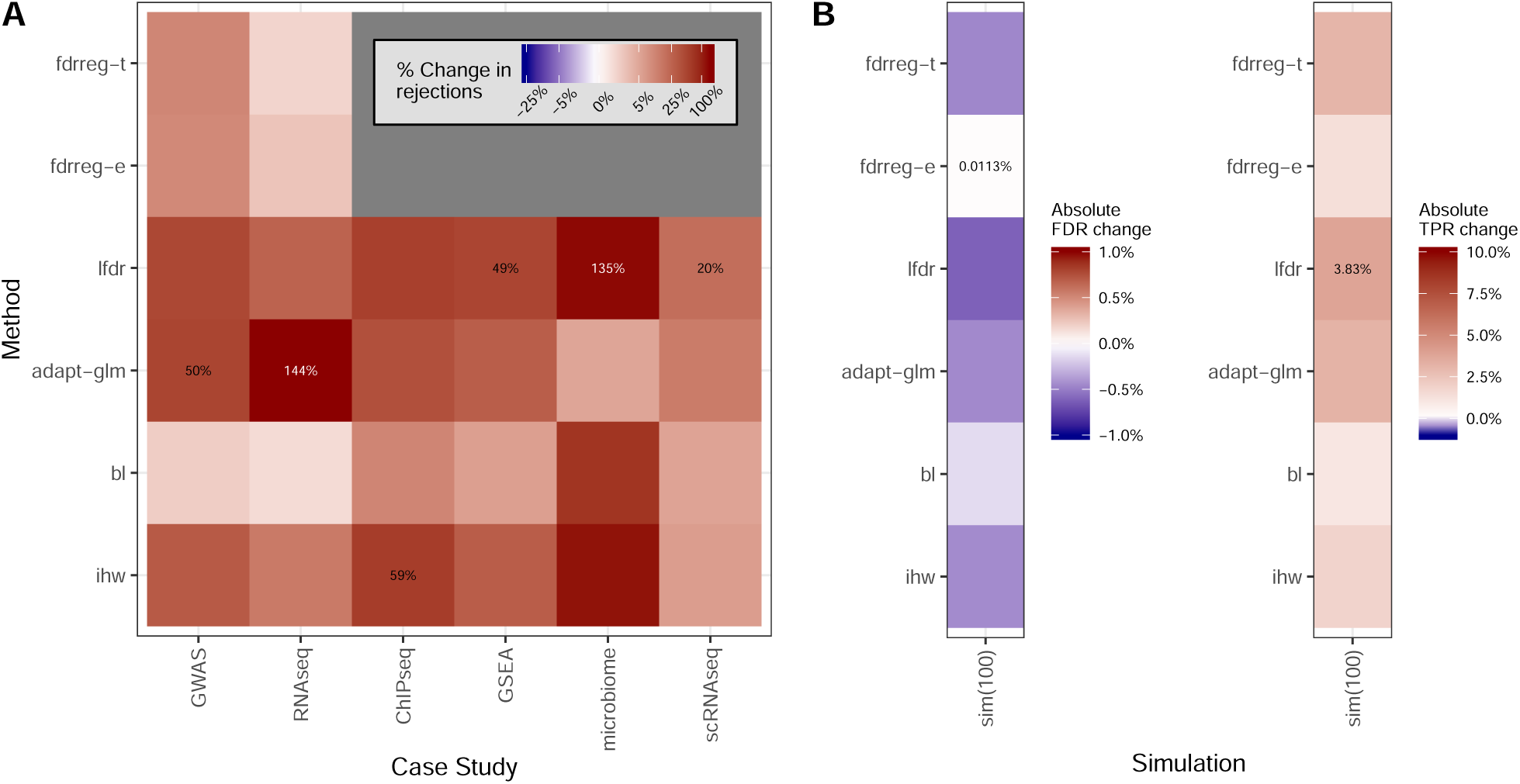
Gain from informative covariate varies by case study. (A) For each method (y-axis) that uses an informative covariate, the percent change in rejections when using an informative covariate as compared to using a completely uninformative covariate is represented by color. This is defined as number of rejections when using the informative covariate divided by the number of rejections when using the uninformative (random) covariate, multiplied by 100. This value is averaged across all datasets and informative covariates used in each case study (x-axis). If rejections were found using the informative covariate but none were found using the uninformative covariate, the percentage was set to 100%. The maximum value of this percentage in each column is labeled. (B) For each method (y-axis) that uses an informative covariate, the absolute percentage change in FDR (left) and TPR (right) in the yeast *in silico* experiments are represented in color (setting with 5 samples in each group, unimodal effect size distribution, and 30% non-null genes). Results are averaged over 100 simulation replications.

**Figure S11.**
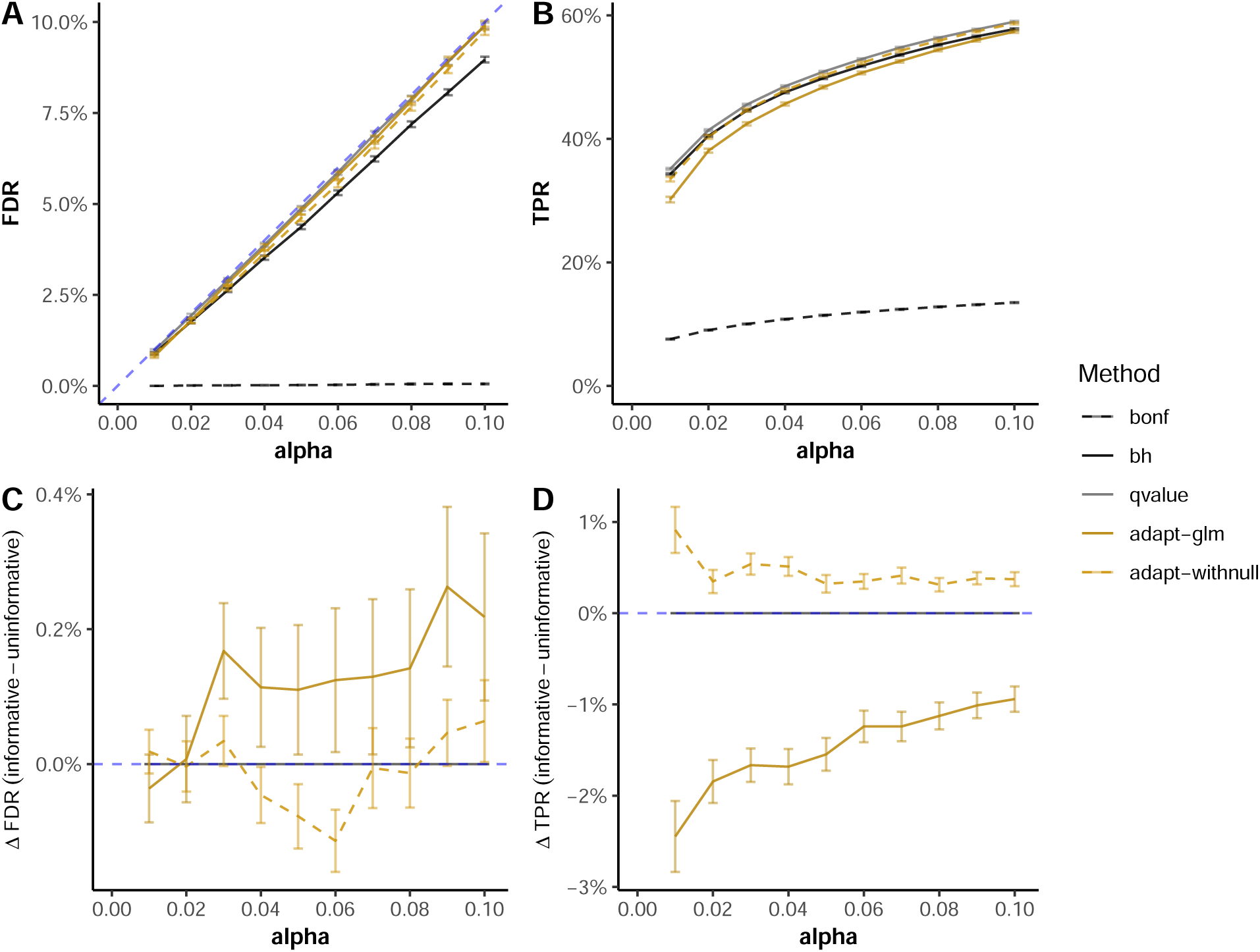
Simulation performance with modified AdaPT parameters. (A) FDR and (B) TPR across nominal FDR thresholds between 0.01 and 0.10 are shown for a subset of methods with the “step” informative covariate (Figure S4). Differences in (C) FDR and (D) TPR between informative and uninformative covariates are also shown. AdaPT with model parameters modified to include a null model is shown as “adapt-withnull” and compared against the default AdaPT call, adapt-glm. Method is denoted by color and the mean value across 100 replications with standard error bars is shown in (A)-(D).

**Figure S12.**
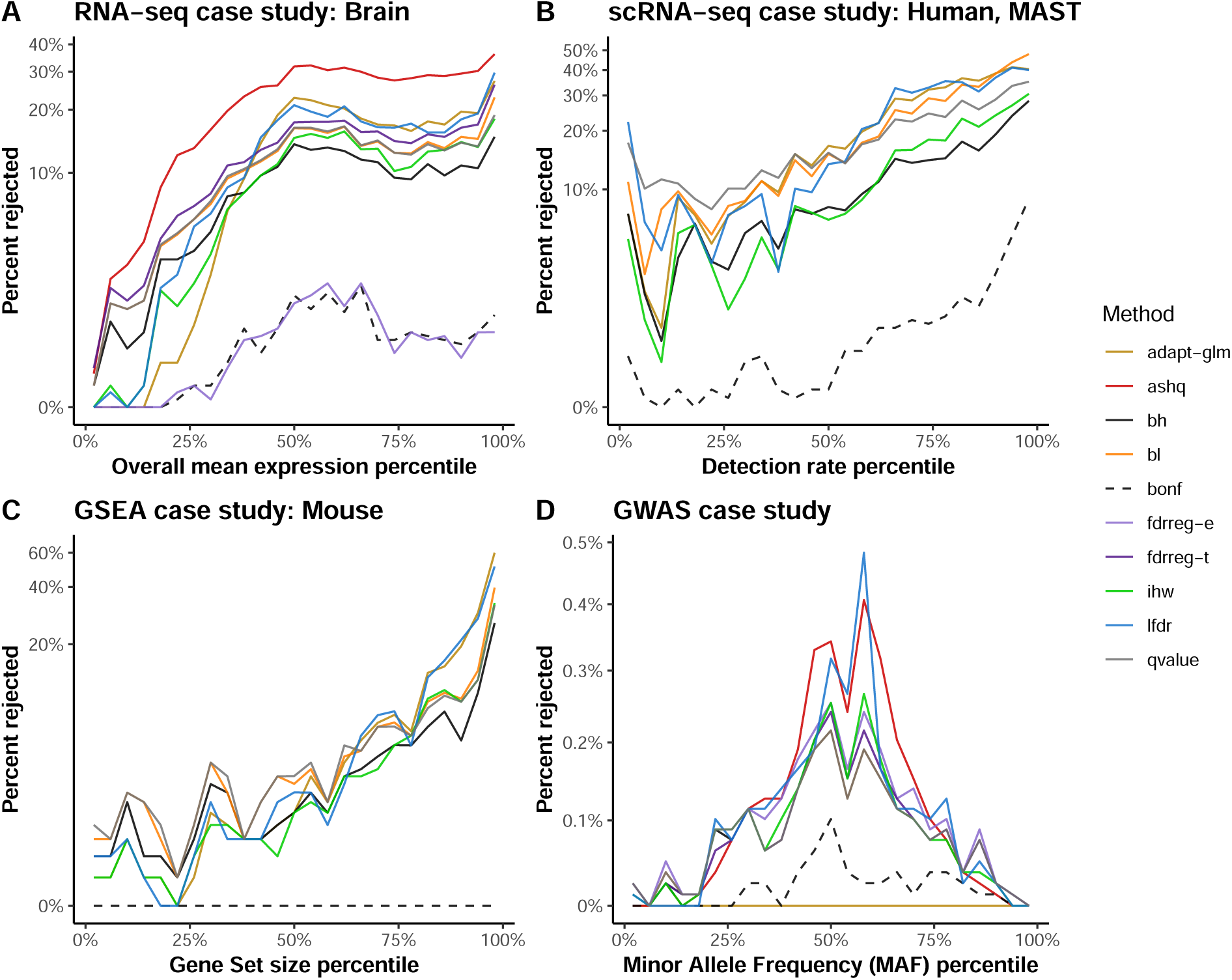
Relationship between informative covariate and rejection rate was highly variable across case studies. Four informative covariates from four different case study datasets were chosen to illustrate the wide variation in the relationship between the informative covariate and the proportion of tests rejected across all FDR controlling methods. (A) Relationship between the overall mean expression percentile and rejection rate in the brain RNA-seq dataset. (B) Relationship between the detection rate percentile and the rejection rate in the Human single cell RNA-seq dataset. (C) Relationship between the gene set size percentile and the rejection rate in the Mouse GSEA dataset. (D) Relationship between the minor allele frequency (MAF) percentile and the rejection rate in the GWAS case study.

**Figure S13.**
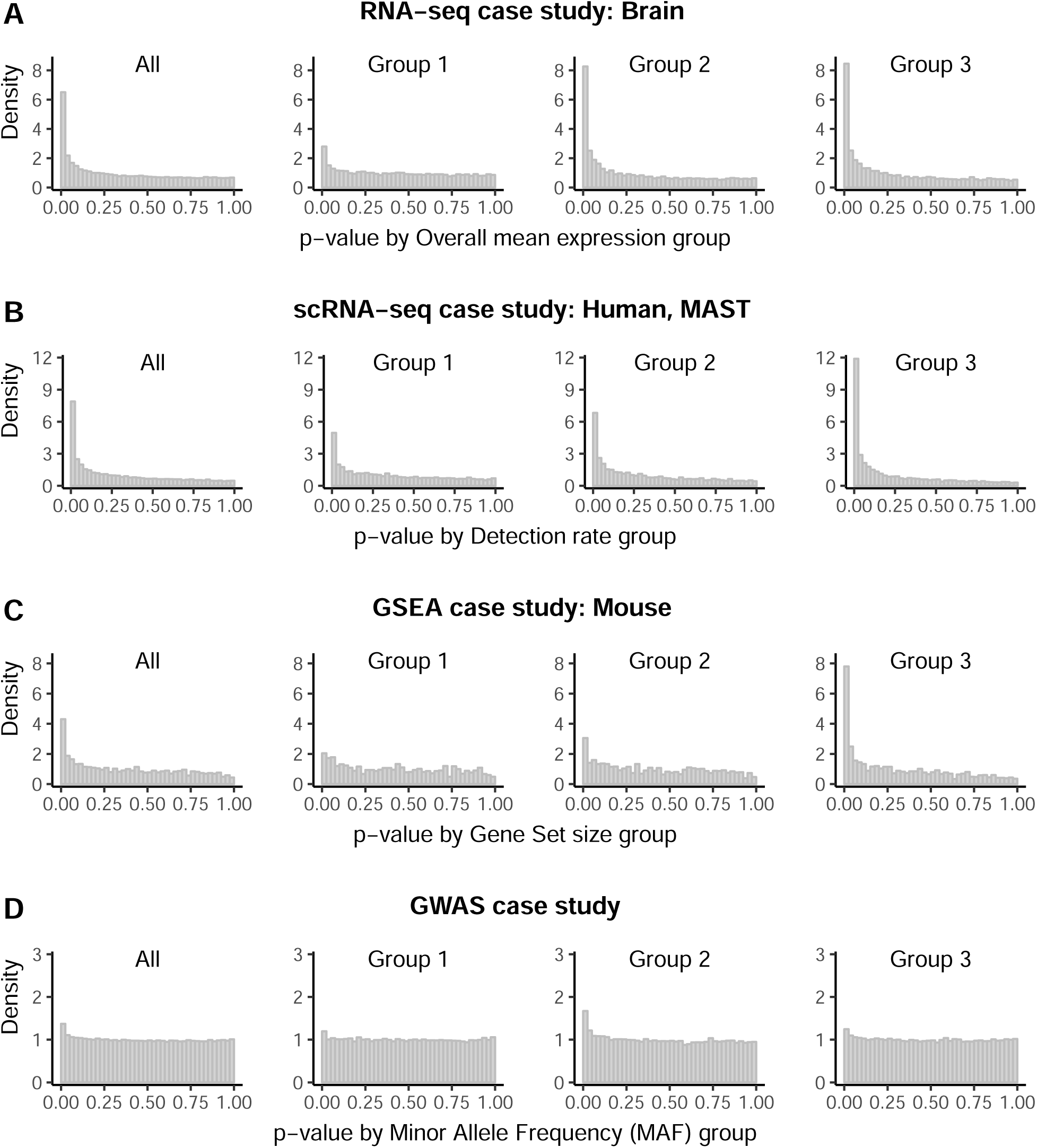
Examples of informative covariates that are independent under the null hypothesis. Distribution of *p*-values overall (left-most panel) and in three approximately equal sized bins by informative covariate (right panels) for (A) the brain dataset in the RNA-seq case study using overall mean expression as the informative covariate, (B) the human dataset in the single-cell RNA-seq case study analyzed by MAST using the detection rate as the informative covariate, (C) the mouse dataset of the GSEA case study using the gene set size as the informative covariate, and (D) the GWAS case study using the minor allele frequency as the informative covariate.

**Figure S14.**
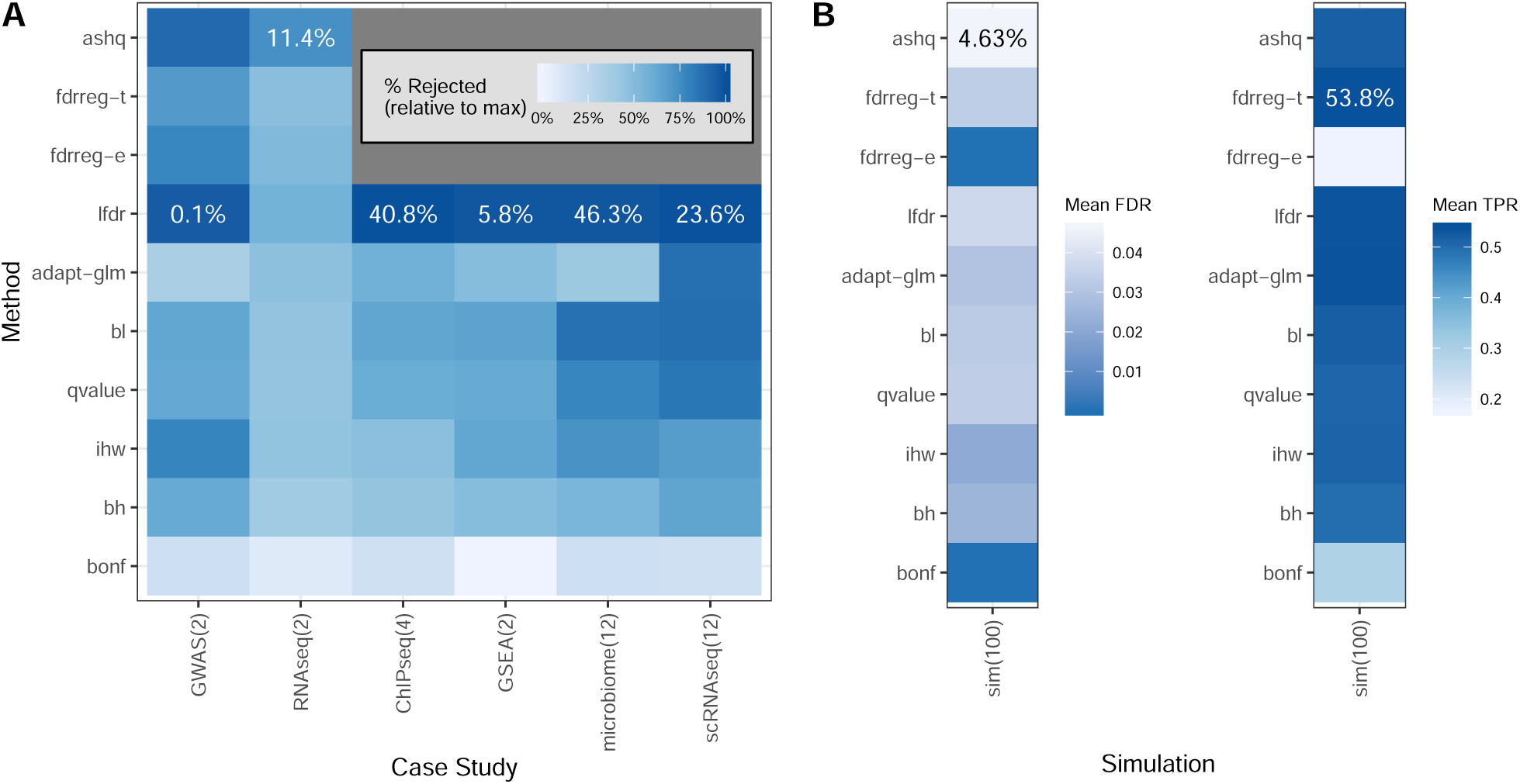
Case study evaluation. (A) Mean proportion of maximum rejections (color) across all datasets and informative covariates used for each case study (column) and FDR correction method (row). In each column, the maximum proportion of rejections out of the total possible number of comparisons is displayed. (B) Mean FDR and TPR (values displayed in each tile and represented by color) in yeast simulation study at target FDR level 0.05 across sample size settings and 100 replications for each FDR correction method (row). Numerical values for the data in (A) are displayed in Additional file 1: Table S4

**Figure S15.**
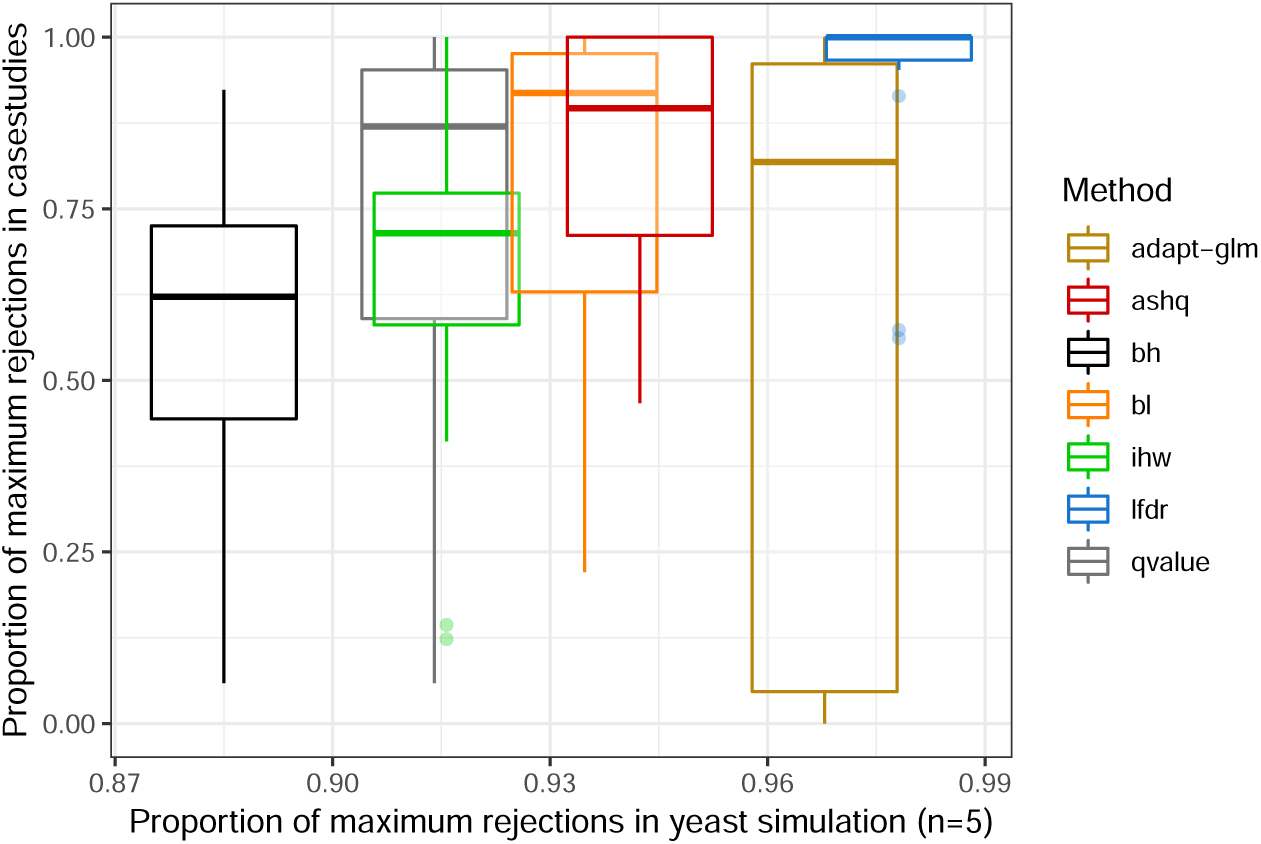
Comparison of the number of rejections in yeast simulation and case studies. Boxplots of the proportion of maximum rejections across all case studies (y-axis) is shown for each method, where the x-axis position reflects the proportion of maximum rejections in the yeast simulation (setting with sample size 5 in each group, unimodal effect size distribution, and 30% non-null genes). The alpha cutoff to determine rejections is 0.05.

**Figure S16.**
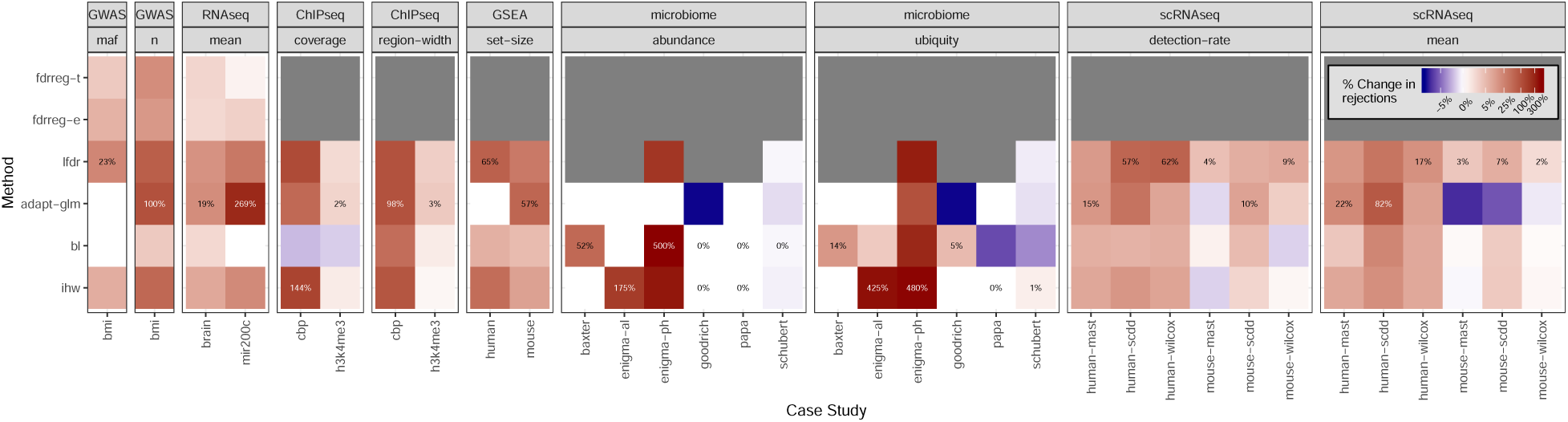
Gain from informative covariate varies by dataset and covariate in case studies. For each method (y-axis) that uses an informative covariate, the percent change in rejections when using an informative covariate as compared to using a completely uninformative covariate is represented by color. This is defined as number of rejections when using the informative covariate divided by the number of rejections when using the uninformative (random) covariate, multiplied by 100. This is shown separately for each dataset (grouped by case study and informative covariate, x-axis). If rejections were found using the informative covariate but none were found using the uninformative covariate, the percentage was set to 100%. The maximum value of this percentage in each column is labeled. The informative covariate used in each case study is listed in Table 1. For case studies with more than one covariate, the covariate is denoted in the column labels

**Table S1.**
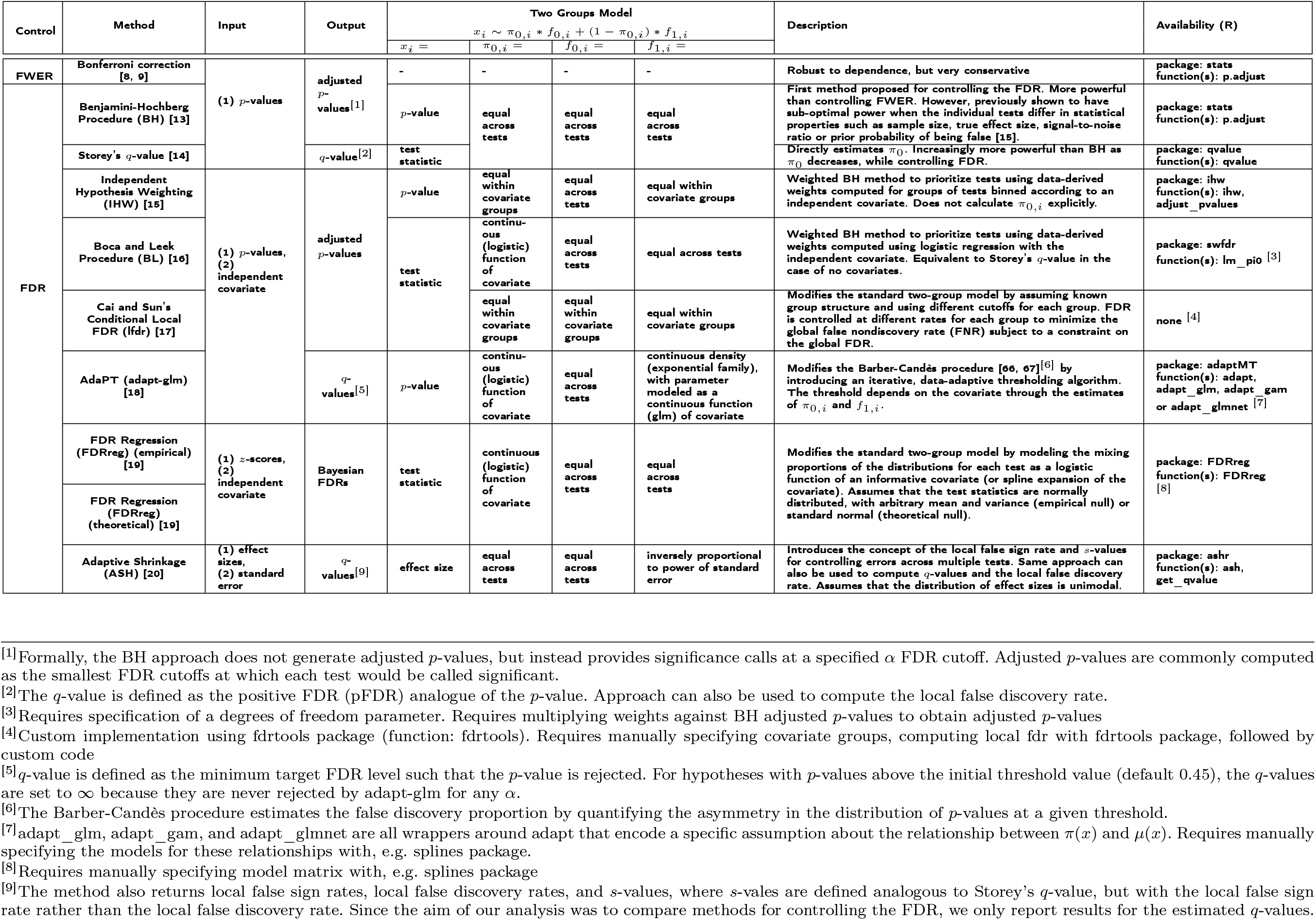
Approaches to adjust for multiple comparisons across hypothesis tests. The family-wise error rate (FWER) is the probability of at least one false discovery. The false discovery rate (FDR) is the expected fraction of false discoveries among all discoveries. FDR adjusted *p*-values are defined as adjusted *p*-values that have control FDR at nominal Type 1 error (*α*) level. π_0_ represents the proportion of null hypothesis tests.

**Table S2.**
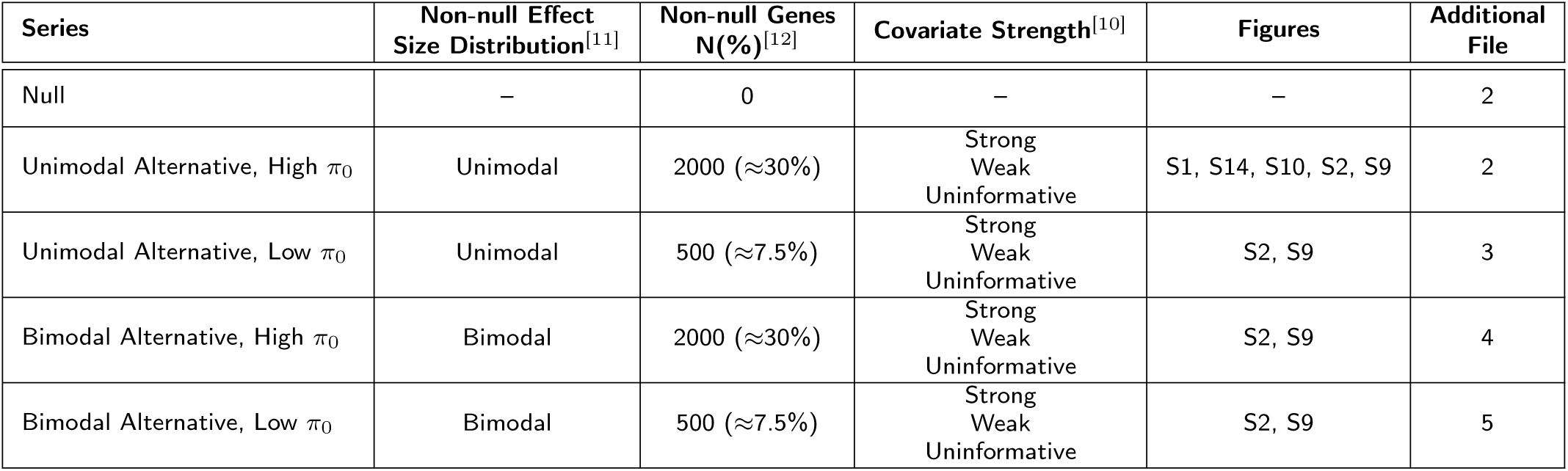
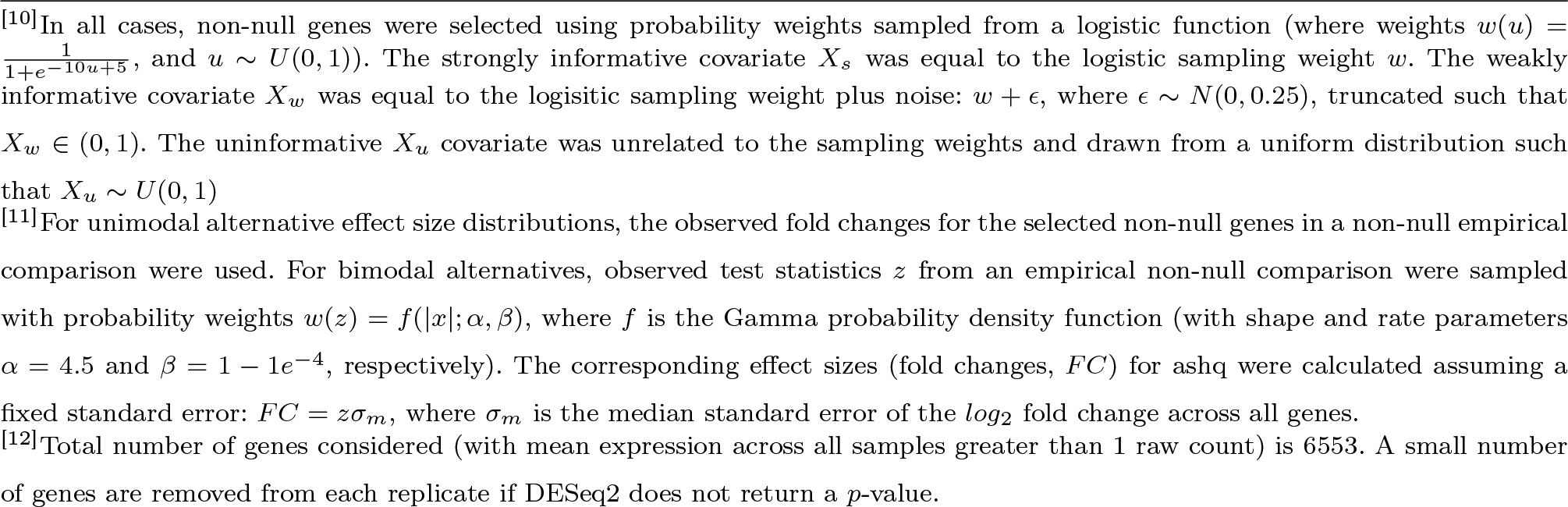
Yeast *in silico* experiment settings. The results from each series of simulations is reported as a separate Additional file in the supplementary materials, with the exception of the Null series, which is combined with the ‘Unimodal Alternative, High π_0_’ series. Both the ‘Null’ and ‘Unimodal Alternative, High π_0_’ series are also evaluated in the Polyester *in silico* experiments, and these results are provided as a separate separate file.

| Series | Non-null Effect Size Distribution<sup>[11]</sup> | Non-null Genes N(%)<sup>[12]</sup> | Covariate Strength<sup>[10]</sup> | Figures | Additional File |
| --- | --- | --- | --- | --- | --- |
| Null | – | 0 | – | – | 2 |
| Unimodal Alternative, High $\pi_0$ | Unimodal | 2000 ($\approx 30\%$) | Strong<br>Weak<br>Uninformative | S1, S14, S10, S2, S9 | 2 |
| Unimodal Alternative, Low $\pi_0$ | Unimodal | 500 ($\approx 7.5\%$) | Strong<br>Weak<br>Uninformative | S2, S9 | 3 |
| Bimodal Alternative, High $\pi_0$ | Bimodal | 2000 ($\approx 30\%$) | Strong<br>Weak<br>Uninformative | S2, S9 | 4 |
| Bimodal Alternative, Low $\pi_0$ | Bimodal | 500 ($\approx 7.5\%$) | Strong<br>Weak<br>Uninformative | S2, S9 | 5 |

<sup>[10]</sup>In all cases, non-null genes were selected using probability weights sampled from a logistic function (where weights $w(u) = \frac{1}{1+e^{-10u+5}}$ , and $u \sim U(0,1)$ ). The strongly informative covariate $X_s$ was equal to the logistic sampling weight $w$ . The weakly informative covariate $X_w$ was equal to the logistic sampling weight plus noise: $w + \epsilon$ , where $\epsilon \sim N(0,0.25)$ , truncated such that $X_w \in (0,1)$ . The uninformative $X_u$ covariate was unrelated to the sampling weights and drawn from a uniform distribution such that $X_u \sim U(0,1)$
<sup>[11]</sup>For unimodal alternative effect size distributions, the observed fold changes for the selected non-null genes in a non-null empirical comparison were used. For bimodal alternatives, observed test statistics $z$ from an empirical non-null comparison were sampled with probability weights $w(z) = f(|z|; \alpha, \beta)$ , where $f$ is the Gamma probability density function (with shape and rate parameters $\alpha = 4.5$ and $\beta = 1 - 1e^{-4}$ , respectively). The corresponding effect sizes (fold changes, $FC$ ) for ashq were calculated assuming a fixed standard error: $FC = z\sigma_m$ , where $\sigma_m$ is the median standard error of the $\log_2$ fold change across all genes.
<sup>[12]</sup>Total number of genes considered (with mean expression across all samples greater than 1 raw count) is 6553. A small number of genes are removed from each replicate if DESeq2 does not return a $p$ -value.

**Table S3.**
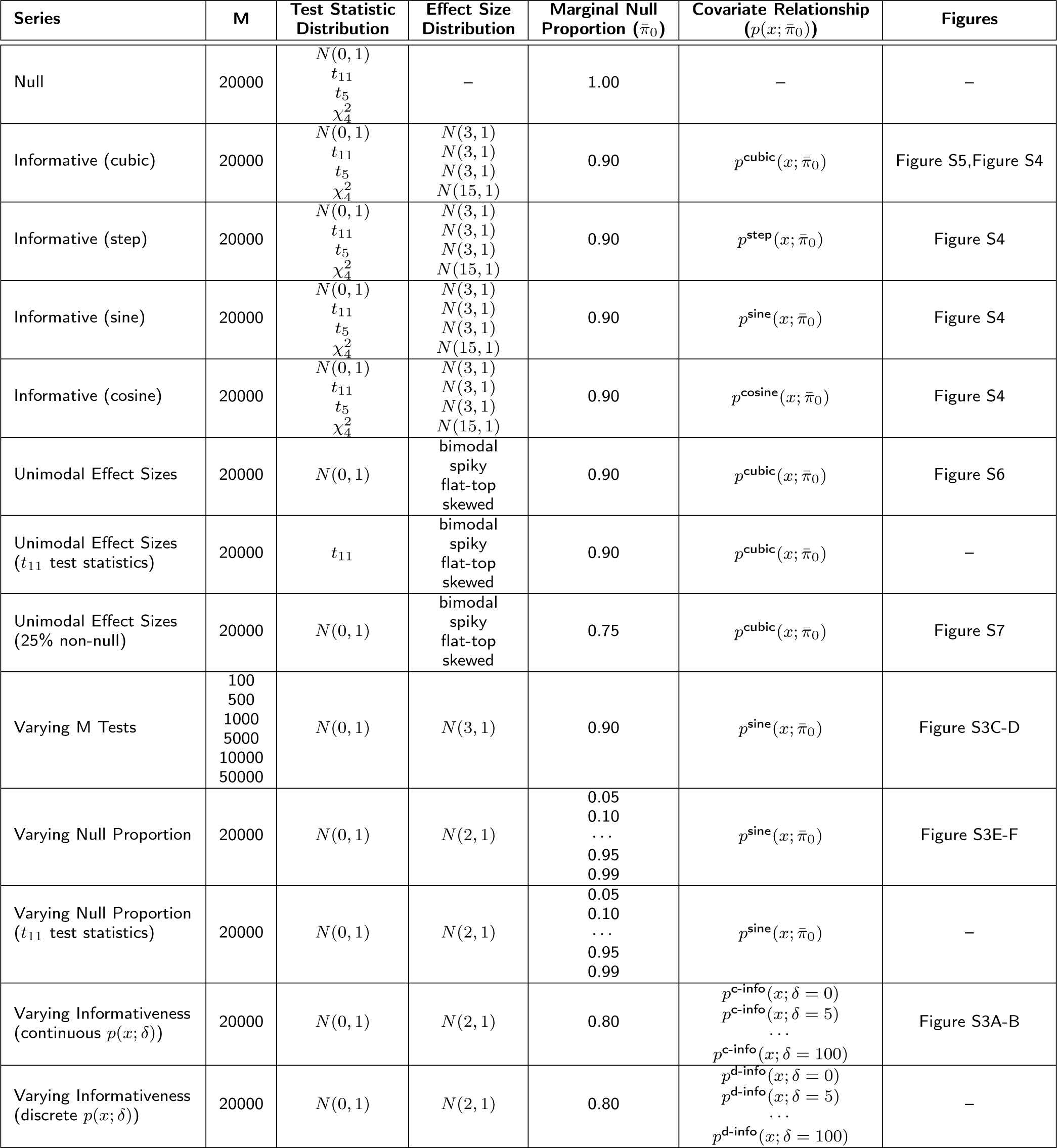
Simulation settings. The results from each series of simulations is reported as a separate Additional file in the supplementary materials.

| Series | M | Test Statistic Distribution | Effect Size Distribution | Marginal Null Proportion ($\bar{\pi}_0$) | Covariate Relationship ($p(x; \bar{\pi}_0)$) | Figures |
| --- | --- | --- | --- | --- | --- | --- |
| Null | 20000 | $N(0, 1)$<br>$t_{11}$<br>$t_5$<br>$\chi^2_4$ | – | 1.00 | – | – |
| Informative (cubic) | 20000 | $N(0, 1)$<br>$t_{11}$<br>$t_5$<br>$\chi^2_4$ | $N(3, 1)$<br>$N(3, 1)$<br>$N(3, 1)$<br>$N(15, 1)$ | 0.90 | $p^{\text{cubic}}(x; \bar{\pi}_0)$ | Figure S5, Figure S4 |
| Informative (step) | 20000 | $N(0, 1)$<br>$t_{11}$<br>$t_5$<br>$\chi^2_4$ | $N(3, 1)$<br>$N(3, 1)$<br>$N(3, 1)$<br>$N(15, 1)$ | 0.90 | $p^{\text{step}}(x; \bar{\pi}_0)$ | Figure S4 |
| Informative (sine) | 20000 | $N(0, 1)$<br>$t_{11}$<br>$t_5$<br>$\chi^2_4$ | $N(3, 1)$<br>$N(3, 1)$<br>$N(3, 1)$<br>$N(15, 1)$ | 0.90 | $p^{\text{sine}}(x; \bar{\pi}_0)$ | Figure S4 |
| Informative (cosine) | 20000 | $N(0, 1)$<br>$t_{11}$<br>$t_5$<br>$\chi^2_4$ | $N(3, 1)$<br>$N(3, 1)$<br>$N(3, 1)$<br>$N(15, 1)$ | 0.90 | $p^{\text{cosine}}(x; \bar{\pi}_0)$ | Figure S4 |
| Unimodal Effect Sizes | 20000 | $N(0, 1)$ | bimodal<br>spiky<br>flat-top<br>skewed | 0.90 | $p^{\text{cubic}}(x; \bar{\pi}_0)$ | Figure S6 |
| Unimodal Effect Sizes ($t_{11}$ test statistics) | 20000 | $t_{11}$ | bimodal<br>spiky<br>flat-top<br>skewed | 0.90 | $p^{\text{cubic}}(x; \bar{\pi}_0)$ | – |
| Unimodal Effect Sizes (25% non-null) | 20000 | $N(0, 1)$ | bimodal<br>spiky<br>flat-top<br>skewed | 0.75 | $p^{\text{cubic}}(x; \bar{\pi}_0)$ | Figure S7 |
| Varying M Tests | 100<br>500<br>1000<br>5000<br>10000<br>50000 | $N(0, 1)$ | $N(3, 1)$ | 0.90 | $p^{\text{sine}}(x; \bar{\pi}_0)$ | Figure S3C-D |
| Varying Null Proportion | 20000 | $N(0, 1)$ | $N(2, 1)$ | 0.05<br>0.10<br>...<br>0.95<br>0.99 | $p^{\text{sine}}(x; \bar{\pi}_0)$ | Figure S3E-F |
| Varying Null Proportion ($t_{11}$ test statistics) | 20000 | $N(0, 1)$ | $N(2, 1)$ | 0.05<br>0.10<br>...<br>0.95<br>0.99 | $p^{\text{sine}}(x; \bar{\pi}_0)$ | – |
| Varying Informativeness (continuous $p(x; \delta)$) | 20000 | $N(0, 1)$ | $N(2, 1)$ | 0.80 | $p^{\text{c-info}}(x; \delta = 0)$<br>$p^{\text{c-info}}(x; \delta = 5)$<br>...<br>$p^{\text{c-info}}(x; \delta = 100)$ | Figure S3A-B |
| Varying Informativeness (discrete $p(x; \delta)$) | 20000 | $N(0, 1)$ | $N(2, 1)$ | 0.80 | $p^{\text{d-info}}(x; \delta = 0)$<br>$p^{\text{d-info}}(x; \delta = 5)$<br>...<br>$p^{\text{d-info}}(x; \delta = 100)$ | – |

**Table S4.** Case Study results. For each case study and method, mean proportion of maximum number of rejections by any method at *α* = 0.05, as shown in the left panel of Figure S14. Range is given in parentheses. A ‘-’ indicates that the method was not applied to the specified case studies.

| Method | GWAS | RNA-seq | ChIP-seq | GSEA | Microbiome | scRNA-seq |
| --- | --- | --- | --- | --- | --- | --- |
| ashq | 0.9 (0.79-1) | 0.73 (0.47-1) | - | - | - | - |
| fdrreg-t | 0.69 (0.66-0.72) | 0.48 (0.42-0.55) | - | - | - | - |
| fdrreg-e | 0.78 (0.71-0.84) | 0.52 (0.04-1) | - | - | - | - |
| lfdr | 0.96 (0.91-1) | 0.57 (0.56-0.57) | 1 (1-1) | 0.98 (0.95-1) | 0.98 (0.96-1) | 1 (0.96-1) |
| adapt-glm | 0.34 (0-0.69) | 0.47 (0.38-0.56) | 0.58 (0.19-0.93) | 0.5 (0-1) | 0.41 (0-1) | 0.87 (0.61-1) |
| bl | 0.64 (0.57-0.7) | 0.45 (0.4-0.49) | 0.64 (0.28-0.92) | 0.65 (0.51-0.79) | 0.78 (0.22-1) | 0.88 (0.63-1) |
| qvalue | 0.63 (0.55-0.7) | 0.44 (0.4-0.48) | 0.61 (0.28-0.92) | 0.61 (0.48-0.75) | 0.7 (0.06-1) | 0.84 (0.59-1) |
| ihw | 0.79 (0.73-0.84) | 0.45 (0.41-0.48) | 0.48 (0.12-0.82) | 0.63 (0.61-0.65) | 0.65 (0.34-0.92) | 0.68 (0.5-0.78) |
| bh | 0.62 (0.54-0.69) | 0.39 (0.38-0.4) | 0.44 (0.06-0.82) | 0.5 (0.44-0.55) | 0.53 (0.04-0.92) | 0.64 (0.45-0.77) |
| bonf | 0.17 (0.15-0.18) | 0.08 (0.04-0.11) | 0.16 (0-0.32) | 0 (0-0) | 0.16 (0-0.46) | 0.16 (0.07-0.23) |

## 1 Supplementary *in silico* experiment results

Here we summarize the benchmarking results of fdrreg-e, which was excluded from the main results due to its unstable and often inferior performance compared to its counterpart fdrreg-t. The difference between these two implementations of FDRreg is that fdrreg-t assumes the null distribution of test statistics is standard normal, while fdrreg-e estimates the null distribution of test statistics empirically. We find this estimation procedure to be sensitive to settings of the distribution of effect sizes and proportion of non-null tests in particular.

### 1.1 Summary of fdrreg-e performance

We found that while modern FDR methods generally led to a higher true positive rate (TPR), or power, in the *in silico* experiments and simulations, fdrreg-e was sometimes as conservative as the Bonferroni correction (Figure S2). The increase in TPR of fdrreg-e sometimes showed substantial improvement over modern methods in several simulation settings (Figures S2, S3, S4, S5, S6, S7). However, these gains were often accompanied by a lack of FDR control, highlighting the sensitivity of fdrreg-e to underlying model assumptions.

#### 1.1.1 Number of tests

We observed that the FDR control of fdrreg-e was sensitive to the number of tests in simulation. Specifically, FDR was substantially inflated when fdrreg-e was applied to fewer than 1,000 tests (Figure S3C). FDR control generally improved as the number of tests increased.

#### 1.1.2 Proportion of non-null tests

The performance of fdrreg-e was particularly sensitive to extreme changes in the proportion of non-null tests. In simulations, fdrreg-e exhibited inflated FDR when the proportion of non-null hypotheses was near 50% (Figure S3E), and suffered from low TPR when there were more than 20% non-null hypotheses, excluding settings where the FDR was not controlled (Figure S3F). In the yeast *in silico* experiments, we also observed that fdrreg-e was more conservative when the proportion of non-null genes was 30% compared to when it was 7.5% (Additional file 1: Figure S2). Similar results were also observed in a series of simulations where unimodal effect sizes were used when the proportion of non-null tests was increased from 10% (Additional file 1: Figure S6) to 25% (Additional file 1: Figure S7).

#### 1.1.3 Distribution of test statistics

We observed that the performance of fdrreg-e declined when the normality assumption of the test statistic was violated (Additional file 1: Figure S5B-C). FDR was considerably inflated when it was applied to *t*-distributed test statistics. As expected, the increase in FDR was greater for the heavier-tailed *t* distribution with fewer degrees of freedom (Additional file 1: Figure S5B).

#### 1.1.4 Distribution of effect sizes

In addition to distributional assumptions on the test statistic, empirical FDRreg requires distributional assumptions on the effect size. Specifically, the empirical null framework used in fdrreg-e relies on [1] to estimate the distribution of null test statistics which requires that all test statistics with values near zero are null, referred to as the ‘zero assumption’. If this is not true, as is the case when the effect sizes are unimodal, the estimation of the null distribution is unidentifiable and may become overly wide, resulting in conservative behavior.

To investigate the sensitivity of these methods to the distribution of effect sizes, multiple distributions of the effect sizes were considered in both yeast *in silico* experiments and simulations. Both unimodal effect size distributions and those following the assumption of fdrreg-e, with most non-null effects greater than zero (Additional file 1: Figure S5A), were considered. While most simulations included the latter, simulations were also performed with a set of unimodal effect size distributions described in [2] (Additional file 1: Figure S6 and S7). In the yeast *in silico* experiments, two conditions were investigated - a unimodal and a bimodal case.

As expected, we observed that when the zero assumption of empirical FDRreg is violated, fdrreg-e was more conservative in both the yeast *in silico* experiments (Additional file 1: Figure S2) and in simulation (Additional file 1: Figures S6 and S7).

We also note that while it is simple to check distributional assumptions on the overall distribution of test statistics or effect sizes, in practice it is impossible to check the distributional assumptions of empirical FDRreg under the alternative, since they rely on knowing which tests are non-null.

### 2 Supplementary case study results

To illustrate what types of covariates may be informative in controlling the FDR in different computational biology contexts, we compared the methods using six case studies including genome-wide association testing (Section 2.1), gene set analysis (Section 2.2), detecting differentially expressed genes in bulk RNA-seq (Section 2.3) and single-cell RNA-seq (Section 2.4), differential binding in ChIP-seq (Section 2.5), and differential abundance testing in the microbiome (Section 2.6). Here we provide additional results for each case study to complement the summary provided in the main text. For full details of the analyses and results, refer to Additional files 21-41.

#### 2.1 Case-study: Genome-Wide Association Studies

Genome-Wide Association Studies (GWAS) are typically carried out on large cohorts of independent subjects in order to test for association of individual genetic variants with a phenotype. The genetic variants are generally measured using microarrays containing probes for up to several million Single Nucleotide Polymorphisms (SNPs). These SNP probes target single base-pair DNA sites that have been shown to vary across a population. To boost power, meta-analyses of GWAS group together many studies, commonly including hundreds of thousands to millions of SNPs, with heterogeneous effect sizes and a wide range of sample size at each loci.

We analyzed a GWAS experiment that carried out a meta-analysis of hundreds of thousands of individuals for more than two million SNPs for association of genetic variants with Body Mass Index (BMI) [3]. As informative covariates, we considered (1) the minor allele frequency (MAF), or the proportion of the population which exhibits the less common allele, and (2) the number of samples for which each SNP was tested for association in the corresponding meta-analysis. In total, 196,969 approximately independent SNPs (out of 2,456,142) were included in the FDR analysis.

For each covariate, we examined whether its rank was associated with the *p*-value distribution. As expected, larger values of the sample size resulted in an enrichment for smaller *p*-values. Additionally, intermediate values of the MAF were associated with an enrichment for smaller *p*-values. This is expected since an MAF near 0.5 balances the number of samples with each allele, thereby maximizing power to detect a difference. For both covariates, the distribution of moderate to large *p*-values appeared uniform and independent of the value of the covariate. For methods that include a covariate, similar numbers of SNPs were rejected at the 0.05 level for either covariate.

For both informative covariates, we found lfdr, ihw, and fdr-e rejected the largest number of hypotheses, followed by fdr-t. The sample size covariate appeared to be more informative than MAF for lfdr and ihw, as both methods rejected more than ashq, whereas ashq found more discoveries than all covariate-aware methods that used MAF. Neither covariate seemed to be very informative for bl, as it did not have much gain over bh or qvalue. adapt-glm was more conservative than Bonferroni with the MAF covariate, but was ranked above bl using the sample size covariate. The overlap among the methods was high, with the largest set sizes containing SNPs rejected by all methods except Bonferonni and/or adapt-glm for both covariate comparisons. The next largest set size was the SNPs rejected by ashq exclusively.

#### 2.2 Case-study: Gene set analyses

Gene set analysis is commonly used to provide insights to results of differential expression analysis. These methods aim to identify gene sets such as Gene Ontology (GO) categories or biological pathways that exhibit a pattern of differential expression. One class of methods, called overrepresentation approaches, test each gene set for a higher number of differentially expressed genes than expected by chance [4]. Another class of methods, called functional class scoring approaches, test each gene set for a coordinated change in expression [4]. While the former operates on a list of differentially expressed genes and does not consider the magnitude or direction of effect, the latter uses information from all genes, and can even detect small coordinated changes across many genes that are not significantly DE individually. We investigated the use of an informative covariate in GOseq [5], an overrepresentation test, as well as Gene Set Enrichment Analysis (GSEA) [6], a functional class scoring approach. Since the sizes of gene sets differ substantially and these size differences translate into differences in power, we hypothesized that multiple-testing correction in gene set analysis would benefit from methods that incorporate information about set sizes.

We used two RNA-seq datasets that investigated gene expression changes (1) between cerebellum and cerebral cortex [7] and (2) upon differentiation of hematopoietic stem cells (HSCs) into multipotent progenitors (MPP) [8]. We obtained 9,853 and 1,336 differentially expressed genes with FDR below 0.10 (using BH) for the human and mouse datasets, respectively. We observed that for both GSEA and GOseq larger gene sets were more likely to have smaller *p*-values than smaller gene sets. Thus, the covariate was informative. In addition, the covariate appeared to be independent under the null hypothesis for GSEA, as evaluated by the histogram of *p*-values stratified by gene set size bins. However, upon evaluation of the stratified histograms of GOseq *p*-values, we observed that the distribution of *p*-values in the larger range was quite different for different covariate bins. This suggests that gene set size is not independent under the null hypothesis for GOseq, so the assumptions of methods which use an independent covariate are violated. Thus, we do not include the GOseq method in the benchmarking study and instead proceed with GSEA *p*-values only.

In this case study, we excluded the methods fdrreg-e, fdrreg-t, and ashq since they require standard errors and test statistics that GSEA does not provide. Overall lfdr, bl, and ihw rejected more hypotheses compared to the other methods. However, the ranking among these methods was not the same between the different datasets. Fort the mouse dataset, lfdr found the most rejections at smaller *α* levels (less than 0.05), but adapt-glm found the most at higher *α* levels. This was followed by BL, qvalue, and then IHW. For the human dataset, lfdr found the most rejections at all *α* levels, followed by IHW and then BL, and adapt-glm was more conservative than BH for almost all *α* levels. As expected, performance using the random (uninformative) covariate of BL and IHW was almost identical to qvalue and BH, respectively. However, the adapt-glm using the uninformative covariate was quite different in the two datasets, with no rejections in the human, and more rejections than any other method in the mouse (at *α* > 0.05).

#### 2.3 Case-study: Differential gene expression in bulk RNA-seq

High-throughput sequencing of mRNA molecules has become the standard for transcriptome profiling. A central analysis task is to determine which genes are deferentially expressed between two biological conditions. Statistical models have been established to address this question including DESeq2 [9] and edgeR [10]. These methods return per gene *p*-values that are further adjusted for multiple testing, typically using the Benjamini-Hochberg procedure.

We assessed the performance of modern FDR methods in the context of differential expression on two RNA-seq datasets. The first dataset consisted of two tissues of 10 individuals from the *GTEx* project and the second dataset consisted of a mouse knockdown experiment of the microRNA *mir200c*. For FDR methods that can use an informative covariate, we used mean expression across samples. We confirmed that this covariate was indeed informative for both datasets.

For the *GTEx* dataset, ashq found more rejections than any other method. At a FDR of 10%, the number of rejections of ashq was more than twice the number of rejections from any of rest of the methods, and the largest gene set was the set of genes found by ashq and no other methods. Following ashq, lfdr, adapt-glm, and fdrreg-t performed similarly. bl found almost the same number of rejections as qvalue, and ihw found slightly more than bh. fdrreg-e was as conservative as Bonferroni. The ranking of the methods based on the number of rejections was consistent across different strata of the covariate.

For the *mir200c* dataset the ranking of the methods was very different compared to the *GTEx* dataset. Here, fdrreg-e found the most rejections by far, and the largest gene set was the set of genes found by fdrreg-e and no other methods. The next highest ranking methods were lfdr, ihw, and ashq, followed by bl, qval, bh, fdrreg-t, and adapt-glm which all performed similarly. For this dataset, the ranking of methods changed substantially across strata of the covariate. For example, among the hypothesis falling between the 50th and 75th percentile of the covariate, lfdr was ranked second (the next highest ranked method after fdrreg-e) but among the hypothesis between the 75th and 100th percentile of the covariate, ashq was ranked second.

#### 2.4 Case-study: Differential gene expression in single-cell RNA-seq

Over the past 5 years, breakthroughs in microfluidics and droplet-based RNA capture technologies have made it possible to sequence the transcriptome of individual cells rather than populations of cells. Quantification of single-cell RNA-seq (scRNA-seq) reads results in a matrix of counts by cells for each sample. The primary applications of scRNA-seq have been in describing cellular heterogeneity in primary tissues, differences in cellular heterogeneity in disease, and discovery of novel cell subpopulations. Differential gene expression of scRNA-seq is used to determine gene sets which distinguish cell populations within the same biological condition, and between cell populations in different samples or conditions.

In this case-study, we examined differences in gene expression in two different biological systems. First, we detected differentially expressed genes between neoplastic glioblastoma cells sampled from a patient’s tumor core with those sampled from nearby peripheral tissue [11]. In addition, we also detected differentially expressed genes between murine macrophage cells that were stimulated to produce an immune response with an unstimulated population [12]. We carried out differential expression analyses using two different methods developed for scRNA-seq: scDD [13] and MAST [14], as well as the Wilcoxon Rank-Sum test.

We examined the mean nonzero expression and detection rate (defined as the proportion of cells expressing a given gene) as potentially informative covariates. For both datasets and all three differential expression methods, we found that mean nonzero expression and detection rate were both informative and approximately independent under the null hypothesis, satisfying the conditions for suitability of inclusion as an informative covariate for controlling FDR. All methods returned more rejections of genes with high nonzero mean and detection rate. They also tended to slightly favor genes with extremely low detection rate.

Across datasets, covariates, and differential expression tests, lfdr usually found the most rejections, followed by bl and adapt-glm. However, at smaller *α* values, adapt-glm was one of the most conservative methods. The ihw and qvalue methods were next, with their rank dependent the dataset and differential expression test used. While a gain in rejections for ihw over bh was apparent in the human dataset, the performance of ihw was very similar to bh in the mouse dataset.

#### 2.5 Case-study: Differential binding in ChIP-seq

ChIP-seq has been widely used to detect protein binding regions and histone modifications in DNA. Testing difference of ChIP-seq signals between conditions usually contains two steps: firstly, defining sets of regions for which the ChIP-seq coverage are quantified; secondly, comparing quantified coverages for testing the statistical significance of differential binding regions. In the first step, regions can be defined by peak calling from samples, based on their signal in sliding windows [15], or by *a priori* interest. In this study we benchmarked results from the latter two approaches by analyzing H3K4me3 data from two widely studied cell lines. Because H3K4me3 is an active marker of gene expression, its signal is most active in promoter regions. This allowed us to pursue an analysis of promoters as regions of interest. We also benchmark the results using the sliding window approach csaw to define *de novo* regions on the H3K4me3 dataset as well as an additional dataset comparing CREB protein binding (CRB) in wild-type versus knock-out mice. We exclude ashq and FDRreg methods from the sliding window analyses since csaw does not provide a standard error or test statistic.

Based on observations that differentially bound peaks tend to have higher coverage, we investigated the use of mean coverage as an informative covariate. In the promoter analysis, the *p*-value histograms showed that high coverage groups are more highly enriched for significant *p*-values different, suggesting mean coverage is an informative covariate. Likewise, we observed that wider windows in the *de novo* analysis tend to have more significant *p*-values. The distribution of *p*-values under the null in both cases appeared approximately uniform.

In the promoter analysis, ashq detected the highest number of differential binding regions by far, followed by lfdr, fdrreg-t, bl, qvalue, and adapt-glm, which all performed similarly. In the sliding window analyses, lfdr rejected the most hypotheses in both datasets, followed by adapt-glm, bl, and qvalue, which performed similarly to one another. In both datasets, the next lowest methods were ihw and bh, where the advantage of ihw only observed in the CBP csaw analysis. Finally, fdrreg-e was more conservative than Bonferroni in the promoter analysis.

#### 2.6 Case-study: Differential abundance testing and correlations in microbiome data analysis

16S rRNA sequencing provides an overview of the microbial community in a given sample, and is a common and accessible way to identify relationships between microbial communities and phenotypes of interest. For example, differential abundance testing is often used to identify bacterial taxa which are enriched or depleted in a disease state, and non-parametric correlations between taxa abundances and phenotypes can be calculated when phenotypes of interest are continuous (e.g. body mass index). However, 16S rRNA datasets are highdimensional, noisy, and sparse, and biological effects can be weak, complicating many statistical analyses and limiting power to detect true associations [16, 17]. Furthermore, environmental samples tend to have many thousands of taxa, which further complicates our ability to identify significant associations

We performed differential abundance tests on the OTU and genus levels for three different datasets from the microbiomeHD database: (1) obesity, where we do not expect a large disease-associated signal [16, 18], (2) inflammatory bowel disease (IBD), which seems to have an intermediate number of disease-associated bacteria [19, 20], and (3) infectious diarrhea (Clostridium difficile (CDI) and non-CDI), where the disease-associated signal is very strong [20, 21]. We also performed Spearman correlation tests between OTU relative abundances and the respective values of three geochemical variables, measured from wells from a contaminated former S-3 waste disposal site in the Bear Creek watershed in Oak Ridge, Tennessee, part of the Department of Energy’s Oak Ridge Field Research Center [22]. The geochemical variables were chosen based on their ability to be predicted by the microbial community in [22]: pH, Al, and SO_4_, where we expect strong, intermediate, and weak associations with the microbial abundances, respectively.

We examined the ubiquity (defined as the proportion of samples with non-zero abundance of each taxa) and mean non-zero abundance of taxa as potentially informative covariates. We found that ubiquity was informative and approximately independent under the null hypothesis, satisfying the conditions for suitability of inclusion as an informative covariate for controlling FDR. Mean non-zero abundance appeared less informative than ubiquity, as it typically showed a less striking pattern in diagnostic plots of *p*-values by covariate value.

OTU-level differential abundance analyses did not have sufficient power to detect any significant differences in the IBD, CRC, and obesity datasets. Similarly, no OTUs correlated with SO_4_ levels and ubiquity was not informative in this case. In addition, very few rejections were found in the CRC dataset at the genus level. Consequently, ubiquity was not informative in these “null” analyses and almost all FDR-correction methods found no significant associations. These “null” results are excluded from the results in the main text.

For the other analyses (genus-level differential abundance, OTU-level differential abundance in diarrhea, OTU-level correlation analyses for pH and Al), ubiquity was informative and the FDR-correction methods which incorporated this information tended to recover more significant associations than naive methods. When there were enough tests for it to be applied, lfdr typically found the most rejections. This was usually followed by bl and qvalue, with the gain of bl over qvalue variable by dataset. In the correlation analyses, however, ihw found more rejections than bl and qvalue. The performance of adapt-glm was usually highly variable, both within a dataset and across datasets: it either had a very different ranking at different *α* levels (obesity), found the among the most rejections (correlation of pH), or found no rejections at all (IBD, CRC). In cases with very few tests (e.g. genus-level analyses), ihw used only 1 covariate bin and reduced to bh as expected.

